# Whole Genome 3D Blood Biopsy Profiling of Canine Cancers: Development and Validation of EpiSwitch Multi-Choice Array-Based Diagnostic Test

**DOI:** 10.1101/2024.05.22.595358

**Authors:** Ewan Hunter, Matthew Salter, Ryan Powell, Ann Dring, Tarun Naithani, Dominik Vugrinec, Kyrylo Shliaiev, Mutaz Issa, Cicely Weston, Abigail Hatton, Abel Gebregzabhar, Jayne Green, Anthony Blum, Thomas Guiel, Sara Fritz, Davis Seelig, Jaime F. Modiano, Alexandre Akoulitchev

## Abstract

Veterinary oncology has a critical need for an accurate, specific, and sensitive non-invasive (blood) biomarker assay to assess multiple canine oncological indications early to better inform therapeutic interventions. Extended from clinical applications in human oncology, here we report on a novel 3D genomics approach to identify systemic blood biomarkers for canine diffuse large B-cell lymphoma (DLBCL), T-zone lymphoma (TZL), hemangiosarcoma (HSA), histiocytic sarcoma, osteosarcoma, and canine malignant melanoma, in a single assay format that encompasses multiple classes and phenotypes of cancer. In the validation of the independent test cohort the 3D whole-genome profiling in peripheral blood demonstrated high sensitivity and specificity for lymphomas and sarcomas as a class, with accuracy >80%; and high sensitivity and specificity for individual indications, with accuracy >89%. This study demonstrates a 3D genomic approach can be used to develop a non-invasive, blood-based test for multiple choice diagnosis of canine oncological indications. The modular EpiSwitch® Specific Canine Blood (EpiSwitch SCB) test promises to help veterinary specialists to diagnose the disease, make more informed treatment decisions, better utilize alternative effective treatments, minimize or avoid unnecessarily toxicity, and efficiently manage costs and resources.

## Background

> *Everything I know I learned from dogs.*
>
> — *-Nora Roberts, “The Search”*

> *“Dogs’ lives are too short. Their only fault, really.”*
>
> — *-Agnes Sligh Turnbull, “The Wedding Bargain”*

Cancer is the predominant cause of death in adult dogs. Similarly to Charles Darwin’s remarks on aspects of our own species’ evolution, this significant disease burden in dogs betrays ‘the indelible stamp of [their] origin’ wherein heavy selection pressures for certain desirable behavioural or phenotypic traits, combined with our success increasing their life expectancy beyond the evolutionarily selected lifespan^1^, unwittingly had as an unintended consequence, an excess of multiple types of cancer observed in this species. In the United States alone, it is estimated that 4.2 to 6 million new cases of cancer are diagnosed each year^2,3^. The lifetime risk of cancer and associated mortality rates in dogs have been estimated to be approximately 30% and have been noted to vary by breed^4^, although recent data suggest that size and biological aging account for most of the observed variation in cancer susceptibility^5,6^. For greater context, non-neoplastic conditions, including those arising from traumatic, infectious, metabolic, inflammatory, degenerative, toxic, congenital, and vascular processes, individually account for ≤10% of adult dog deaths^7^. When accounting for age, sex, and weight, the lifetime incidence of cancer in purebred and mixed-breed dogs is reported to be comparable^8,9^.

Malignant lymphomas are one of the most common cancers in dogs, with an estimated incidence of 25 per 100,000^10^. While they may occur at any age, they are predominately diagnosed in older dogs^11,12^. Of these, the most frequently encountered subtype is diffuse large B-cell lymphoma (DLBCL), accounting for 40% of all canine lymphomas and approximately 70% of canine B-cell lymphomas^13^

Marginal zone lymphoma (MZL) is less common but the expansion of immunophenotyping and molecular clonality assessments available to veterinary pathologists are making greater scrutiny possible^14^. Indolent lymphomas make up approximately 29% of all canine lymphomas^15,16^. T-zone lymphoma (TZL) is the most frequently diagnosed indolent lymphoma in dogs^15,16^. Estimations speaking to TZL incidence specifically vary widely, with two publications’ estimations ranging from 15.5 and 62%^17,18^. Histopathology and immunohistochemistry (IHC) remain the clinical gold standard for diagnosing and classifying lymphomas^18,19^

It has been reported that approximately 7% of all malignant tumours in dogs are melanocytic tumours ^20^. Oral melanocytic tumours make up 30-40% of all canine oral neoplasms^21^. Regardless of their location, malignant melanocytic neoplasms are diagnosed via cytology and/or histopathology ^22^. The sensitivity and specificity of this diagnostic method are excellent for pigmented melanocytic neoplasms but can drop significantly with diagnostic attempts of amelanotic melanocytic neoplasms ^23–25^. If the diagnosis of amelanotic melanocytic neoplasms remains evasive with microscopic evaluation, IHC for melanocyte-specific markers is required for definitive confirmation^26–30^.

Sarcomas comprise approximately 10-15% of malignant tumours in dogs. Of these, bone is the primary anatomic location of origin in 20% of cases, while the remaining 80% are found in soft tissue ^31^. Hemangiosarcoma (HSA) is among the more commonly occurring soft tissue sarcomas with an incidence of 2.8-5.0% of all diagnosed cancers, with primary sites of occurrence in the spleen, heart, liver, and skin^31^. Diagnosis of cutaneous HSA is performed via analysis of histopathology from incisional or excisional biopsies and may require IHC for definitive diagnosis owing to its heterogeneous makeup^32–34^. Splenic and liver HSA typically requires abdominal ultrasonography and three-view chest radiography. Definitive diagnosis requires histopathological examination and differential IHC is recommended^32^. Cardiac HSA diagnosis is made by fluid cytology but may not always be achieved due to the intrinsic challenges associated with sample acquisition, low cellular exfoliation and elevated haemodilution^32,35–37^.

Histiocytic sarcoma represents approximately 4% of all canine cancers, originating from antigen-presenting cells with primary sites in the lymph nodes, kidneys, liver, and central nervous system ^31^. Cutaneous histiocytic sarcoma can be diagnosed with relative ease by cytopathologic examination but as with all subtypes of histiocytic sarcoma, IHC staining is required to provide a definitive diagnosis^38^.

Within non-soft tissue sarcomas, osteosarcoma is the most common bone cancer in dogs, occurring in the appendicular (∼85%) and axial skeleton^31,39,40^. Definitive diagnosis of osteosarcoma requires histological observation of osteoid production by malignant osteoblast cells^31^.

Securing a definitive diagnosis and classification of cancer in dogs falls on a spectrum of varying degrees of difficulty as location, subtypes, complicating local conditions, and specific biopsy requirements are factored in. Given the invasive nature of tissue biopsies requiring sedation or anaesthesia, the economic costs associated with the process, and the specialized technical training required to achieve an accurate and definitive classification, several alternative diagnostic methods are gradually gaining favour^41^. For example, fine needle aspiration alone or in conjunction with immunocytochemistry, flow cytometry, and/or polymerase chain reaction (PCR) for antigen receptor rearrangement (PARR) are among the most widely employed contemporary alternatives used to diagnose canine lymphomas^42^. Several liquid biopsy tools have been reported to aid in the diagnosis and subtyping of canine lymphoma, in addition to facilitating early disease detection, minimal residual disease monitoring, and the capacity to guide therapy^43–46^.

Unfortunately, many dogs with a presumptive diagnosis of malignancy are euthanized without a definitive diagnosis, underscoring the need for reliable and affordable tests that can better inform treatment options and prognosis for dogs with cancer. Emerging liquid biopsy and molecular testing platforms are addressing this unmet need^47^. Here, we report on the first evidence of a novel liquid biopsy biomarker modality focused on the changes in 3-dimensional genomic architecture (3D genomics) in peripheral blood cells – already utilized in DLBCL and other oncological applications in humans^19,48–51^ – to establish the presence of the canine cancers described above, as well as to report on cancer indications in training. Exploration of higher-order genomic architecture has revealed that the 3D configuration of chromatin plays a critical role as an epigenetic regulator of gene expression in pathological and non-pathological phenotypes^52,53^.

Of central importance are 3D chromosomal conformations signatures (CCSs) derived from combinations of long-range DNA contacts that establish a regulatory fragmentation of the active genome, linking together genetic risks and epigenetic oversight of genome regulation^54^. By determining long-range chromosome interactions, *i.e.*, chromatin loops, at particular loci via chromosome conformation capture (3C) technologies, one can derive a biomarker that provides an instructive view of the regulatory architecture possessed by the 3D genome^55^. The novel biomarker *EpiSwitch®* platform is grounded on 3C and has reduced to practice and meets the regulatory requirements of a clinical biomarker assay. The heart of the platform is the unification of high-throughput, high-resolution screening with the *EpiSwitch* Explorer microarray platform, machine learning algorithms, testing on independent validation cohorts, and a PCR-based format for stratifying biomarker signature^19,49,51,56–58^. This well-established phenomenon of CCSs being the multi-omics integrator and regulatory interface of the cell has been validated in the *EpiSwitch* platform in an extensive range of indications including detection of disease (pre-symptomatic and symptomatic), prognosis, and predictive response to treatment^19,48–51,56,59–65^. Two of the *EpiSwitch* blood-based biomarker applications are practiced today in US and UK as reimbursable clinical tests: a Checkpoint Inhibitor Response Test (CiRT) for prediction of treatment response in immuno-oncology^48^, and a prostate cancer screening test (PSE)^50^.

A salient and increasingly appreciated aspect of the regulatory architecture inherent in the 3D genome is that the heritable information in DNA far exceeds the linear genetic and epigenetic histone modifications that academic literature has classically explored. These traditional linear genetic formats and epigenetic histone modifications are in fact “encoded” in 3D genomic folding and architecture, constituting part of the epigenetic memory, reproducing itself with high fidelity through innumerable cell divisions, and are intimately related to cell metabolic and epigenetic states of the cell. More simply, the 3D genome is the storage media of the heritable footprint-genetic, epigenetic, and metabolic - of the cellular network regulation.

Visualizing how a whole blood biopsy could, by utilizing the EpiSwitch microarray platform and machine learning, provide diagnostic and prognostic insights into such extensive pathologies and processes, it is crucial to understand the extent to which individual chromosome conformations could be synchronized in the context of the multicellular organism and all the cross talk between the cells. Exosome traffic fosters the active exchange of epigenetic factors between the cells. These extracellular vesicles, with their cargoes of metabolites, nucleic acids, non-coding RNA, lipids, and peptides transfer horizontally from cells of origin to recipient cells and result in the modulation of immune cells (representing systemic changes), as well as modulation of secondary sites. Together this generates systemic epigenetic synchronization and changes in 3D genomic profiles, detectable by the *EpiSwitch* microarray platform^66–69^. Here we have deployed the stratifying capabilities of the whole genome *EpiSwitch* 3D genomic array profiling based on peripheral blood biopsy to several prevalent canine cancers: lymphomas – DLBCL and TZL; HSA, histiocytic sarcoma, osteosarcoma; and canine malignant melanoma.

Given the challenge of a multi-choice outcome, we have developed an approach based exclusively on array readouts. The two-step classifier identifies first the strong systemic network signatures for lymphomas, sarcomas, and melanomas, shared by each class, and then identifies individual indication within the class. With batch alignment and internal controls, the performance of the array-based stratifications was then evaluated in validation cohorts for accuracy and specificity against other cancer indications.

## Material and Methods

### Samples

Canine whole blood samples, 150 in total, were imported under licenses ITIMP19.0336 and ITIMP22.0063 (Animal & Plant Health Agency) from the Animal Cancer Care and Research Program, University of Minnesota, St. Paul, MN, USA. Samples represented healthy controls and cases of DLBCL, TZL, HSA, histiocytic sarcoma, osteosarcoma, and canine malignant melanoma (Supplemental Table 1). When available, the annotations also include the age and breed of the individual dogs. All samples were profiled on EpiSwitch Canine Whole-genome 3D Explorer Array. Samples were either used in *EpiSwitch* screening and discovery stage, or in validation evaluation. Those samples were not used in any stages of the classifier model development.

### Preparation of 3D genomic templates

*EpiSwitch* 3D libraries, chromosome conformation analytes converted to sequence-based tags, were prepared from frozen whole blood samples. Using *EpiSwitch* protocols following the manufacturer’s instructions for *EpiSwitch* Explorer Array kits (Oxford BioDynamics Plc), samples were processed on the Freedom EVO 200 robotic platform (Tecan Group Ltd). Briefly, aliquots of 50 µl of whole blood were diluted and fixed with an EpiSwitch buffer containing formaldehyde. Density cushion centrifugation was used to purify intact, fixed nuclei. Following a short detergent-based step to permeabilise the nuclei, restriction enzyme digestion and proximity ligation were used to generate the 3D libraries. Samples were centrifuged to pellet the intact nuclei before purification with an adapted protocol from the QIAmp DNA FFPE Tissue kit (Qiagen) Eluting in 1x TE buffer pH7.5. 3D libraries were quantified using the Quanti-T™ Picogreen dsDNA Assay kit (Invitrogen) and normalised to 5 ng/ml prior to interrogation by PCR.

### Array design

Custom microarrays were designed using the *EpiSwitch* pattern recognition algorithm, which operates on Bayesian-modelling and provides a probabilistic score that a region is involved in long-range chromatin interactions. The algorithm was used to annotate the CanFam 3.1 canine genome assembly across ∼1.1 million sites with the potential to form long-range chromosome conformations^48,62^. The most probable interactions were identified and filtered on probabilistic score and proximity to protein, long non-coding RNA, or microRNA coding sequences. Predicted interactions were limited to *EpiSwitch* sites greater than 10 kb and less than 300 kb apart. Repeat masking and sequence analysis was used to ensure unique marker sequences for each interaction. The *EpiSwitch* Explorer array (Agilent Technologies, Product Code 087165), containing 60-mer oligonucleotide probes was designed to interrogate potential 3D genomic interactions. In total, 964,631 experimental probes and 2,500 control probes were added to a 1 x 1 M CGH microarray slide design. The experimental probes were placed on the design in singlicate with the controls in groups of 250. The control probes consisted of six different *EpiSwitch* interactions that are generated during the extraction processes and used for monitoring library quality. A further four external inline control probe designs were added to detect non-human (*Arabidopsis thaliana*) spike in DNA added during the sample labelling protocol to provide a standard curve and control for labelling. The external spike DNA consists of 400 bp ssDNA fragments from genomic regions of *A. thaliana*. Array-based comparisons were performed described previously, with the modification of only one sample being hybridised to each array slide in the Cy3 channel^62^.

### Statistical analysis

The cohorts of analysed samples were normalised by background correction and quantile normalisation, using the *EpiSwitch* R analytic package, which is built on the Limma Rank Product, tidyverse libraries. The datasets were combined into sample sets by processing batch. Data were corrected for batch effects using ComBat R script. Parametric (Limma R library, Linear Regression) and non-parametric (*EpiSwitch* RankProd R library) statistical methods were performed to identify 3D genomic changes that demonstrated a difference in abundance between classes.

The resulting data from both procedures were further filtered based on p-value and abundance scores (AS). Only 3D genomic markers with p-value <=0.01 and AS (−1.2<; >1.2) were selected. Both filtered lists from Limma and RankProd analysis were compared and the intersection of the two lists was selected for further processing.

### Machine learning and modelling

All analysis for this study was performed using libraries which are developed for the R Statistical Language (R version 4.2.0). Feature engineering of the EpiSwitch Markers was performed using Recursive Feature Elimination (RFE) utilising Xgbtree, The XGBoost algorithm model^70^ was used for final test optimisation. The grid search algorithm was used to optimize the hyper-parameters and learning rate in each iteration. For drawing inferences, we used SHapley Additive exPlanations (SHAP) values that are computed by a game theoretical approach which quantifies the contribution of each feature within a model to the final prediction of an observation. SHAP values were used to reduce the feature space for the cancer specific models^71^.

### Genomic mapping

The 3D genomic markers from the statistically filtered list with the greatest and lowest abundance scores were selected for genome mapping. Mapping was carried out using Bedtools closest function for the 3 closest protein coding loci – upstream, downstream and within the long-range chromosome interaction (Gencode v33). All markers were visualized using the *EpiSwitch* Analytical Portal.

### Mapping to STRING database

The closest protein coding loci for the chromosome interactions found in this study, where inputted to the STRING DB^72^, utilizing the default settings in order to search STRING. The resultant protein-protein interaction data was exported and then imported and visualised in Cytoscape (v3.10.0).

## Results

### Identification of the top predictive 3D genomic markers common for lymphomas as a class

Following the established methodology for *EpiSwitch* array marker analysis, we evaluated systemic marker leads shared by two types of lymphomas – DLBCL and TZL. From nearly 40 million data points across the canine genome, screening of 20 healthy controls vs 20 lymphomas, represented by DBCL and TZL, identified 37 strong systemic EpiSwitch biomarkers. Those biomarkers were then used in array-based stratification on a validation cohort of 35 dogs, representing healthy controls and both lymphomas. Stratification calls were made for presence of lymphomas vs healthy controls as a class (Figure 1 and Supplemental Table 2).

**Figure 1.**
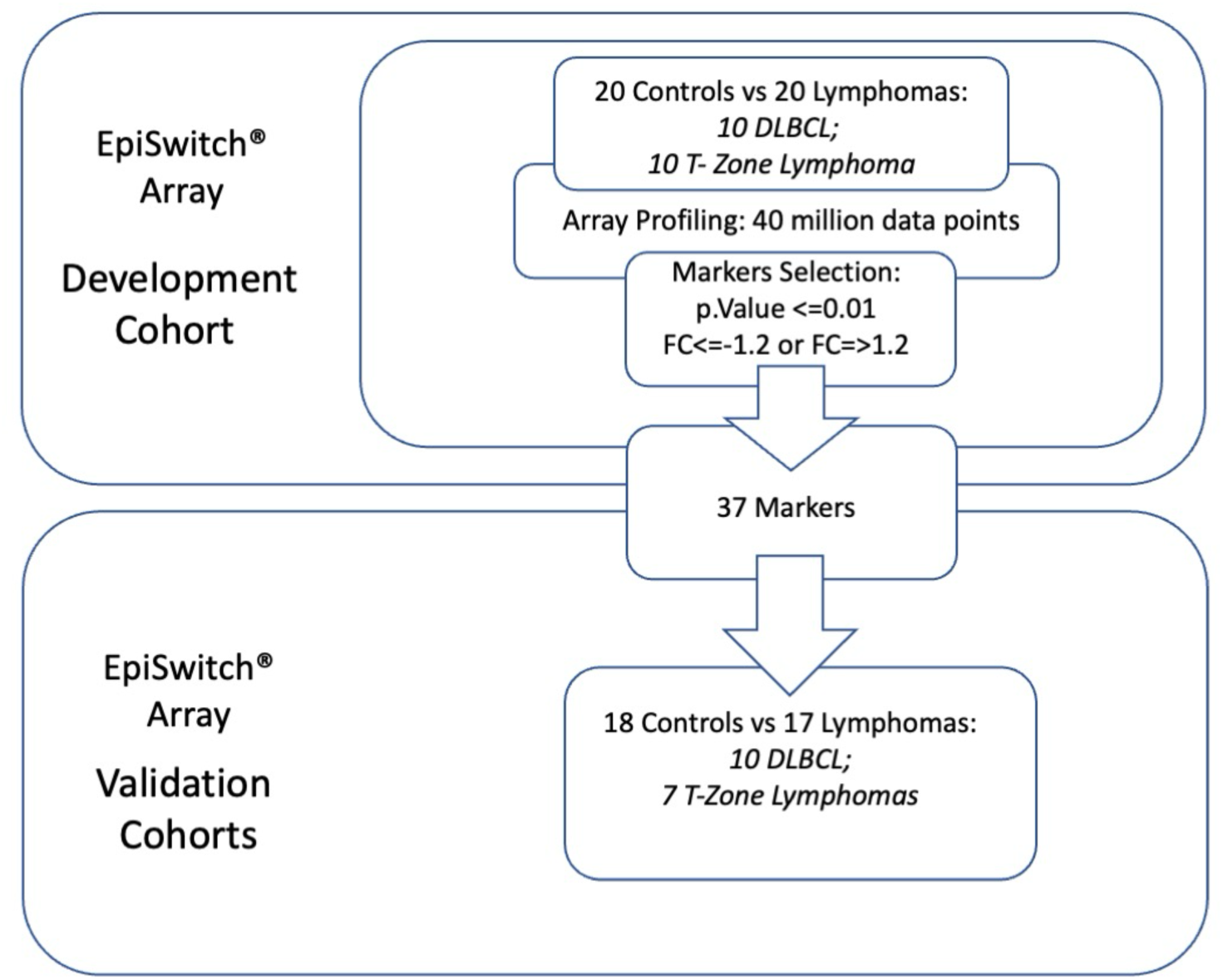
Workflow for development and testing of the 3D genomic classifier model for presence of lymphomas in canine blood. From EpiSwitch array screening profiles accounting for over 40 million data points, 37 top array markers were selected based on p value <=0.01 and fold change (FC) less than −1.2 or more than 1.2. 37 markers qualified through selection were used in array-based stratification for lymphoma on an independent cohort of 35 dogs.

### Testing of the predictive 3D genomic biomarker panel for lymphomas as a class on the independent sample cohort

To access the predictive power of the classifier model, the 37-marker 3D array classifier was validated on an independent test cohort. No samples from that cohort were used in marker selection and building of the model. The *EpiSwitch* platform readouts for the classifier model were uploaded to the *EpiSwitch* Analytical Portal for analysis. Veterinary diagnostic assessment for the test cohort included 18 healthy control samples and a mixture of lymphomas, including 10 DLBCL and 7 (TZL). *EpiSwitch* classifier model calls based on 37-marker model demonstrated high performance of 83% balanced accuracy and 87% positive predictive value in identifying dogs with lymphomas as a class against healthy controls (Table 1).

**Table 1.** Performance of the EpiSwitch array-based biomarker classifier for calling presence of lymphoma as a class, based on common systemic markers shared by DLBCL and TZL across 1 million data point profiles of the 3D genomic architecture. Confusion matrix and test performance statistics for the 37-marker classifier on the 35 dogs in the test cohort.

| Lymphomas as a Class |  |  |  |  |  |
| --- | --- | --- | --- | --- | --- |
| Test | Yes | n | No | n | Total |
| Yes | True Positive | 13 | False Positive | 2 | 15 |
| No | False Negative | 4 | True Negative | 16 | 20 |
| Total |  | 17 |  | 18 |  |

***Table 1. Performance of the EpiSwitch array-based biomarker classifier for calling presence of lymphoma as a class, based on common systemic markers shared by DLBCL and TZL across 1 million data point profiles of the 3D genomic architecture.**
| Results Statistics | Value | 95% CI |
| --- | --- | --- |
| Sensitivity | 76.47% | 50.10% to 93.19% |
| Specificity | 88.89% | 65.29% to 98.62% |
| Positive Likelihood Ratio | 6.88 | 1.81 to 26.10 |
| Negative Likelihood Ratio | 0.26 | 0.11 to 0.63 |
| Disease prevalence | 48.57% | 31.38% to 66.01% |
| Positive Predictive Value | 86.67% | 63.15% to 96.10% |
| Negative Predictive Value | 80.00% | 62.57% to 90.54% |
| Accuracy | 82.68% | 66.35% to 93.44% |

### Identification of the top predictive 3D genomic markers common for sarcomas as a class

Following the established methodology for *EpiSwitch* array marker analysis, we also evaluated systemic marker leads shared by three types of sarcomas – HSA, histiocytic sarcoma, osteosarcoma. From nearly 50 million data points across the canine genome after screening of 20 healthy controls vs 30 sarcomas, represented by all three types, we identified 100 strong systemic *EpiSwitch* biomarkers. Those biomarkers were used in array-based stratification on a validation cohort of 40 dogs, representing healthy controls and all three sarcomas. Stratification calls were made for presence of sarcomas as a class vs healthy control (Figure 2 and Supplemental Table 2).

**Figure 2.**
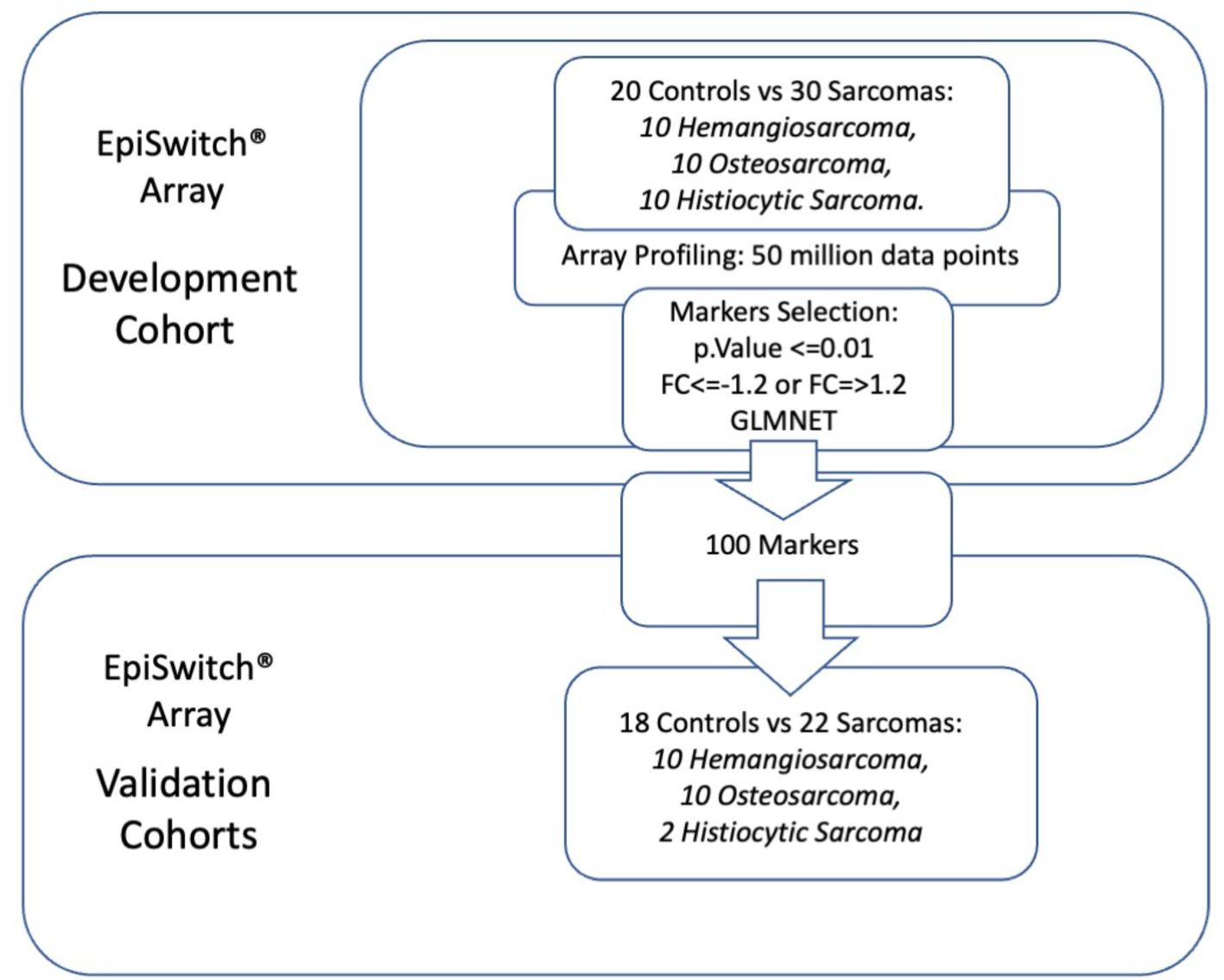
Workflow for development and testing of the 3D genomic classifier model for presence of sarcomas in canine blood. From EpiSwitch array screening profiles accounting for over 50 million data points, 100 top array markers were selected based on p value <=0.01 and fold change (FC) less than −1.2 or more than 1.2, and GLMNET. 100 markers qualified through selection were used in array-based stratification for lymphoma on independent cohort of 40 dogs.

### Testing of the predictive 3D genomic biomarker panel for sarcomas as a class on the independent sample cohort

To access the predictive power of the classifier model, the 100-marker 3D array classifier was validated on an independent test cohort. No samples from that cohort were used in marker selection and building of the model. The *EpiSwitch* platform readouts for the classifier model were uploaded to the EpiSwitch Analytical Portal for analysis. Veterinary diagnostic assessment for the test cohort included 18 healthy control samples and a mixture of HSA, histiocytic sarcoma, osteosarcoma. *EpiSwitch* classifier model calls based on 8-marker model demonstrated high performance of 83% balanced accuracy and 83% positive predictive value in identifying dogs with sarcomas (Table 2).

**Table 2.** Performance of the EpiSwitch array-based biomarker classifier for calling presence of sarcomas as a class, based on common systemic markers shared by all three sarcomas across 1 million data point profiles of the 3D genomic architecture. Confusion matrix and test performance statistics for the 100-marker classifier on the 40 dogs in the test cohort.

| Sarcomas as a Class |  |  |  |  |  |
| --- | --- | --- | --- | --- | --- |
| Test | Yes | n | No | n | Total |
| Yes | True Positive | 19 | False Positive | 4 | 23 |
| No | False Negative | 3 | True Negative | 14 | 17 |
| Total |  | 22 |  | 18 |  |

***Table 2. Performance of the EpiSwitch array-based biomarker classifier for calling presence of sarcomas as a class, based on common systemic markers shared by all three sarcomas across 1 million data point profiles of the 3D genomic architecture.**
| Results Statistics | Value | 95%CI |
| --- | --- | --- |
| Sensitivity | 86.36% | 65.09% to 97.09% |
| Specificity | 77.78% | 52.36% to 93.59% |
| Positive Likelihood Ratio | 3.89 | 1.61 to 9.37 |
| Negative Likelihood Ratio | 0.18 | 0.06 to 0.52 |
| Disease prevalence | 55.00% | 38.49% to 70.74% |
| Positive Predictive Value | 82.61% | 66.33% to 91.97% |
| Negative Predictive Value | 82.35% | 61.31% to 93.22% |
| Accuracy | 82.50% | 67.22% to 92.66% |

### Training and Testing Individual Multi-Choice Classifiers

Having observed strong systemic signatures shared by the indications representing lymphomas and sarcomas as distinct classes according to systemic *EpiSwitch* profiling, we proceeded with two-step classifier models pursuing individual indications within each of the groups.

For that purpose, our approach was to use once again the development cohort of samples for all indications (Figure 3). We have systematically identified top 100 markers from the comparison of each indication to healthy control samples, based on p value and FC, as described earlier. From the total pool we followed only the markers unique for each indication. This has been considered an important filtering step to ensure the high specificity of the multi-choice stratification of individual cancer types. Having ranked the markers, we then pursued only the disease positive markers, *i.e.*, chromosome conformations present for detection in the given indication. Such positive detection markers are associated with negative FC in our data analysis. As the final step, we have conducted further feature reduction of the markers, based on the SHAP plots that identified markers with highest impact. In this analysis we also have processed samples representing melanoma, as a separate indication and a separate class. We then tested a validation cohort of 56 samples, including 18 healthy controls, with the two-step classification, based first on a class call and then, using indication unique markers, with the individual indication calls (Figure 3).

**Figure 3.**
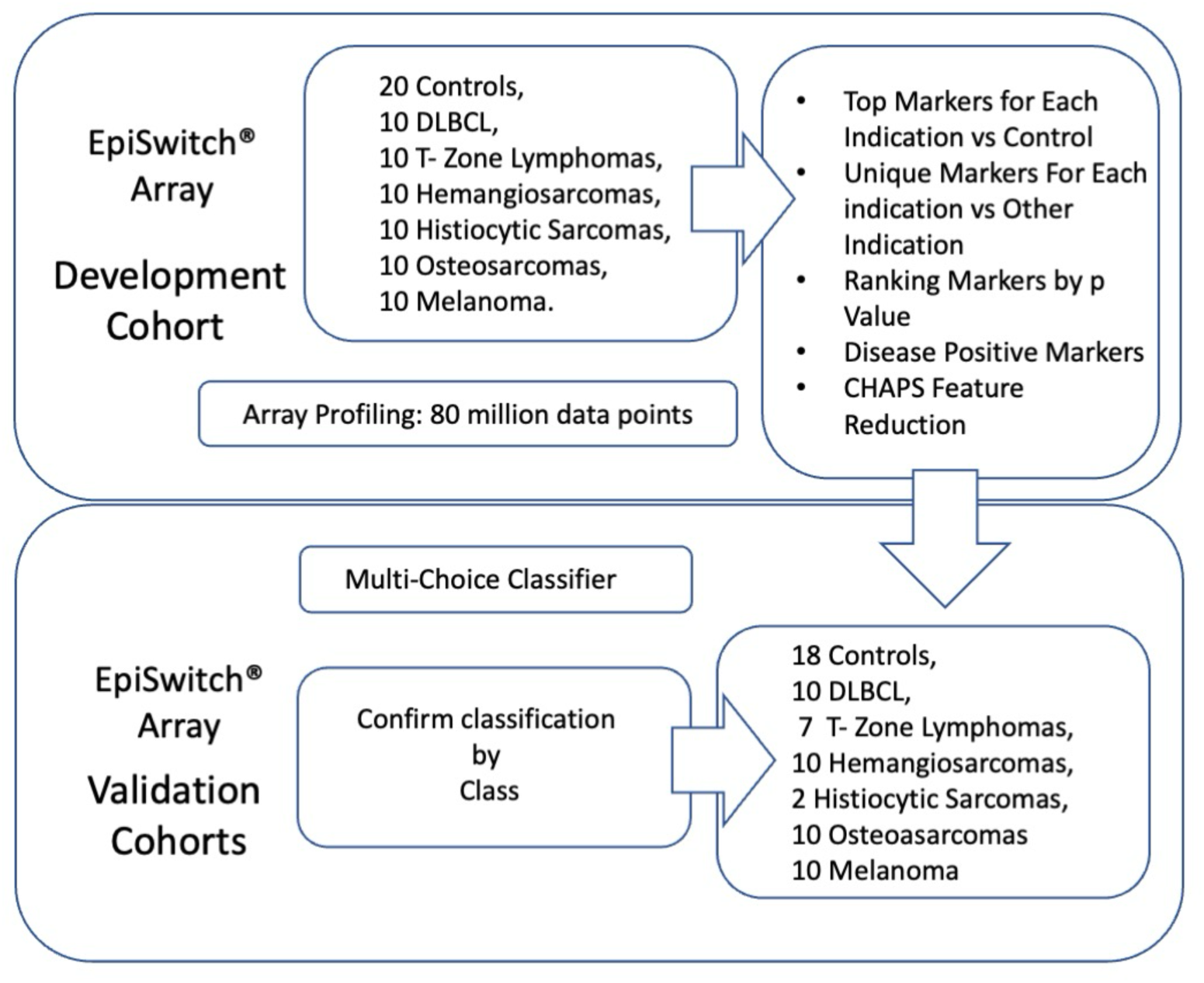
Development of the EpiSwitch array-based biomarker classifier for individual indications. Markers from over 80 million data point were selected with unique profiles for each of the indications within the class and evaluated in multi-choice stratification against healthy controls and other cancer indications.

By using validation cohort of 56 mixed indications and healthy controls, the multi-choice calls for DLBCL demonstrated high positive predictive value of 87.5%, with accuracy against healthy controls at 85.7% and accuracy against all the indications included in the validation cohort at 92.8% (Table 3). Top markers used in the DLBCL individual classifier are listed in the Supplemental Table 2.

**Table 3.** Performance of the EpiSwitch array-based biomarker classifier for calling presence of DLBCL in the validation cohort. Confusion matrix and test performance statistics for the multi-choice classifier against healthy controls (A) and the full validation cohort of 56 samples, including 2 lymphomas, three sarcomas and melanoma.

| A |  |  |  |  |  | B |  |  |  |  |  |
| --- | --- | --- | --- | --- | --- | --- | --- | --- | --- | --- | --- |
| DLBCL vs Healthy Controls |  |  |  |  |  | DLBCL vs All |  |  |  |  |  |
| Test | Yes | n | No | n | Total | Test | Yes | n | No | n | Total |
| Yes | True Positive | 7 | False Positive | 1 | 8 | Yes | True Positive | 7 | False Positive | 1 | 8 |
| No | False Negative | 3 | True Negative | 17 | 20 | No | False Negative | 3 | True Negative | 45 | 48 |
| Total |  | 10 |  | 18 |  | Total |  | 10 |  | 46 |  |

**Table 3. Performance of the EpiSwitch array-based biomarker classifier for calling presence of DLBCL in the validation cohort.**
| Results Statistics | Value | 95%CI |
| --- | --- | --- |
| Sensitivity | 70.00% | 34.75% to 93.33% |
| Specificity | 94.44% | 72.71% to 99.86% |
| Positive Likelihood Ratio | 12.6 | 1.80 to 88.34 |
| Negative Likelihood Ratio | 0.32 | 0.12 to 0.82 |
| Disease prevalence | 35.71% | 18.64% to 55.93% |
| Positive Predictive Value | 87.50% | 49.96% to 98.00% |
| Negative Predictive Value | 85.00% | 68.59% to 93.63% |
| Accuracy | 85.71% | 67.33% to 95.97% |
| Sensitivity | 70.00% | 34.75% to 93.33% |
| Specificity | 97.83% | 88.47% to 99.94% |
| Positive Likelihood Ratio | 32.2 | 4.44 to 233.35 |
| Negative Likelihood Ratio | 0.31 | 0.12 to 0.79 |
| Disease prevalence | 17.86% | 8.91% to 30.40% |
| Positive Predictive Value | 87.50% | 49.13% to 98.07% |
| Negative Predictive Value | 93.75% | 85.32% to 97.48% |
| Accuracy | 92.86% | 82.71% to 98.02% |

The multi-choice calls for TZL demonstrated positive predictive value of 71.43%, with accuracy against healthy controls at 84% and accuracy against all the indications included in the validation cohort at 92.8% (Table 4). Top markers used in the TZL individual classifier are listed in Supplemental Table 2. Given the nature of TZL, we are currently investigating if among the false positives among the healthy control, depending on the breed and age, there might have been pre-symptomatic, undiagnosed true positives.

**Table 4.** Performance of the EpiSwitch array-based biomarker classifier for calling presence of TZL in the validation cohort. Confusion matrix and test performance statistics for the multi-choice classifier against healthy controls (A) and the full validation cohort of 56 samples, including 2 lymphomas, threes sarcomas and melanoma.

| **A** |  |  |  |  |  | **B** |  |  |  |  |  |
| --- | --- | --- | --- | --- | --- | --- | --- | --- | --- | --- | --- |
| TZ vs Healthy Controls |  |  |  |  |  | TZ vs All |  |  |  |  |  |
| Test | Yes | n | No | n | Total | Test | Yes | n | No | n | Total |
| Yes | True Positive | 5 | False Positive | 2 | 7 | Yes | True Positive | 5 | False Positive | 2 | 7 |
| No | False Negative | 2 | True Negative | 16 | 18 | No | False Negative | 2 | True Negative | 47 | 49 |
| Total |  | 7 |  | 18 |  | Total |  | 7 |  | 49 |  |

**Table 4. Performance of the EpiSwitch array-based biomarker classifier for calling presence of TZL in the validation cohort.**
| Results Statistics | Value | 95%CI |
| --- | --- | --- |
| Sensitivity | 71.43% | 29.04% to 96.33% |
| Specificity | 88.89% | 65.29% to 98.62% |
| Positive Likelihood Ratio | 6.43 | 1.60 to 25.76 |
| Negative Likelihood Ratio | 0.32 | 0.10 to 1.05 |
| Disease prevalence | 28.00% | 12.07% to 49.39% |
| Positive Predictive Value | 71.43% | 38.42% to 90.92% |
| Negative Predictive Value | 88.89% | 71.03% to 96.31% |
| Accuracy | 84.00% | 63.92% to 95.46% |
| Sensitivity | 71.43% | 29.04% to 96.33% |
| Specificity | 95.92% | 86.02% to 99.50% |
| Positive Likelihood Ratio | 17.5 | 4.16 to 73.56 |
| Negative Likelihood Ratio | 0.30 | 0.09 to 0.96 |
| Disease prevalence | 12.50% | 5.18% to 24.07% |
| Positive Predictive Value | 71.43% | 37.29% to 91.31% |
| Negative Predictive Value | 95.92% | 87.91% to 98.70% |
| Accuracy | 92.86% | 82.71% to 98.02% |

The multi-choice calls for HSA demonstrated high positive predictive with no false positives within the validation cohort of 56 samples, with accuracy against healthy controls at 85.7% and accuracy against all the indications included in the validation cohort at 92.8% (Table 5). Top markers used in the HSA individual classifier are listed in Supplemental Table 2.

**Table 5.**
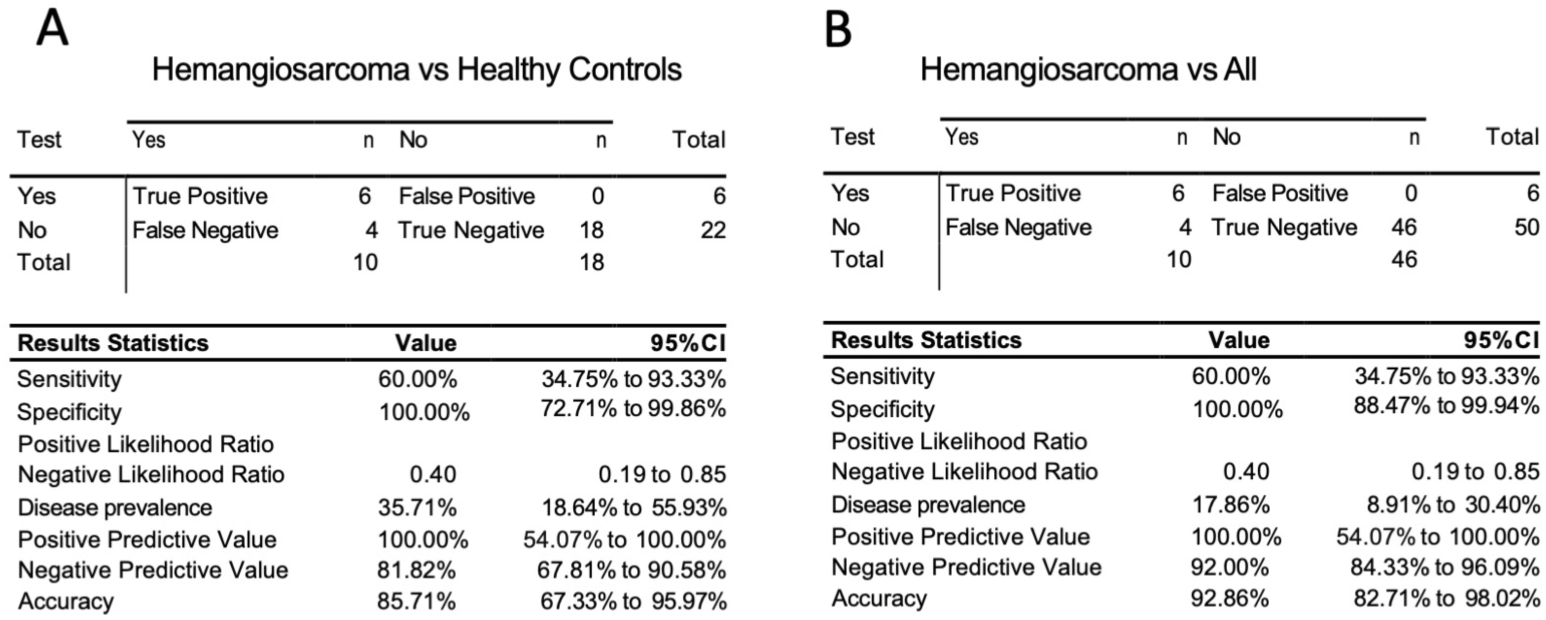
Performance of the EpiSwitch array-based biomarker classifier for calling presence of HSA in the validation cohort. Confusion matrix and test performance statistics for the multi-choice classifier against healthy controls (A) and the full validation cohort of 56 samples, including 2 lymphomas, threes sarcomas and melanoma.

The multi-choice calls for histiocytic sarcoma was underrepresented in the validation cohort, with only 2 histiocytic sarcoma independent samples available within 56 sample validation cohort (Table 6). Interesting to note, that the classifier has demonstrated high specificity, with no false positive calls. Also, the one false negative call on a histiocytic sarcoma sample by the individual classifier still was flagged as a sarcoma class sample at the first stage classification. Top markers used in the histiocytic sarcoma individual classifier are listed in Supplemental Table 2.

**Table 6.**
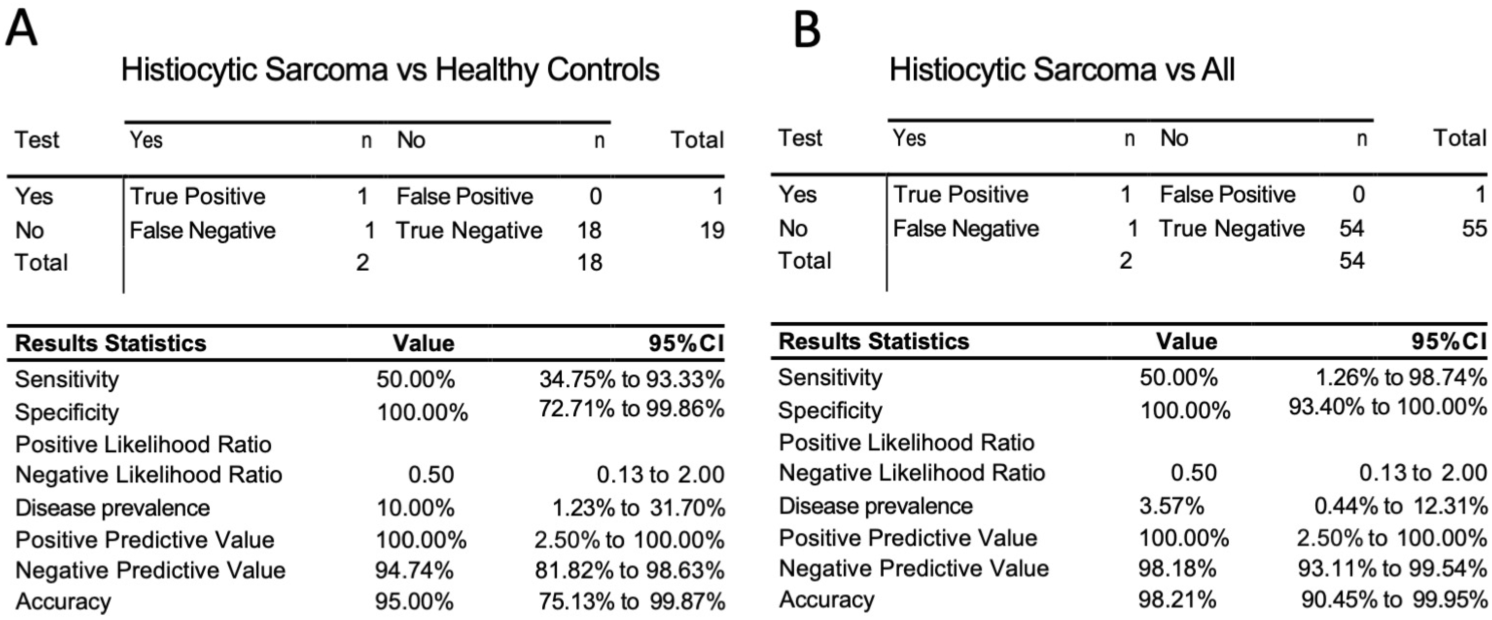
Performance of the EpiSwitch array-based biomarker classifier for calling presence of histiocytic sarcoma in the validation cohort. Confusion matrix and test performance statistics for the multi-choice classifier against healthy controls (A) and the full validation cohort of 56 samples, including 2 lymphomas, threes sarcomas and melanoma.

| Test | Yes | n | No | n | Total |
| --- | --- | --- | --- | --- | --- |
| Yes | True Positive | 1 | False Positive | 0 | 1 |
| No | False Negative | 1 | True Negative | 18 | 19 |
| Total |  | 2 |  | 18 |  |

| Results Statistics | Value | 95%CI |
| --- | --- | --- |
| Sensitivity | 50.00% | 34.75% to 93.33% |
| Specificity | 100.00% | 72.71% to 99.86% |
| Positive Likelihood Ratio |  |  |
| Negative Likelihood Ratio | 0.50 | 0.13 to 2.00 |
| Disease prevalence | 10.00% | 1.23% to 31.70% |
| Positive Predictive Value | 100.00% | 2.50% to 100.00% |
| Negative Predictive Value | 94.74% | 81.82% to 98.63% |
| Accuracy | 95.00% | 75.13% to 99.87% |

B Histiocytic Sarcoma vs All
| Test | Yes | n | No | n | Total |
| --- | --- | --- | --- | --- | --- |
| Yes | True Positive | 1 | False Positive | 0 | 1 |
| No | False Negative | 1 | True Negative | 54 | 55 |
| Total |  | 2 |  | 54 |  |

**Table 6. Performance of the EpiSwitch array-based biomarker classifier for calling presence of histiocytic sarcoma in the validation cohort.**
| Results Statistics | Value | 95%CI |
| --- | --- | --- |
| Sensitivity | 50.00% | 1.26% to 98.74% |
| Specificity | 100.00% | 93.40% to 100.00% |
| Positive Likelihood Ratio |  |  |
| Negative Likelihood Ratio | 0.50 | 0.13 to 2.00 |
| Disease prevalence | 3.57% | 0.44% to 12.31% |
| Positive Predictive Value | 100.00% | 2.50% to 100.00% |
| Negative Predictive Value | 98.18% | 93.11% to 99.54% |
| Accuracy | 98.21% | 90.45% to 99.95% |

The multi-choice calls for osteosarcoma demonstrated high specificity of 80%, positive predictive value of 66.7%, negative predictive value of 66.67%, with accuracy against healthy controls at 78.57% and accuracy against all the indications included in the validation cohort at 89.29% (Table 7). Top markers used in the osteosarcoma individual classifier are listed in Supplemental Table 2. Given the nature of osteosarcoma, we are also currently investigating if among the false positives among the healthy controls, depending on the breed and age, there might have been pre-symptomatic, undiagnosed true positives.

**Table 7.**
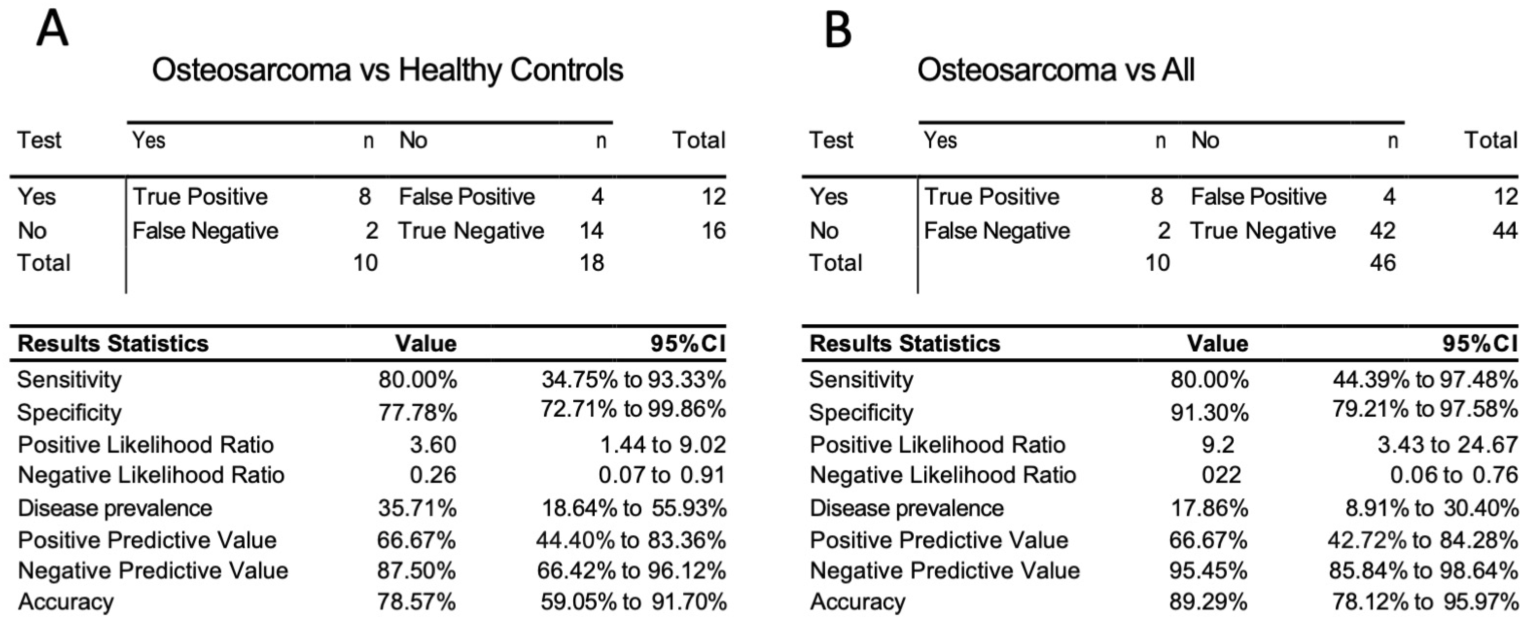
Performance of the EpiSwitch array-based biomarker classifier for calling presence of osteosarcoma in the validation cohort. Confusion matrix and test performance statistics for the multi-choice classifier against healthy controls (A) and the full validation cohort of 56 samples, including 2 lymphomas, threes sarcomas and melanoma.

| Test | Yes | n | No | n | Total |
| --- | --- | --- | --- | --- | --- |
| Yes | True Positive | 8 | False Positive | 4 | 12 |
| No | False Negative | 2 | True Negative | 14 | 16 |
| Total |  | 10 |  | 18 |  |

| Results Statistics | Value | 95%CI |
| --- | --- | --- |
| Sensitivity | 80.00% | 34.75% to 93.33% |
| Specificity | 77.78% | 72.71% to 99.86% |
| Positive Likelihood Ratio | 3.60 | 1.44 to 9.02 |
| Negative Likelihood Ratio | 0.26 | 0.07 to 0.91 |
| Disease prevalence | 35.71% | 18.64% to 55.93% |
| Positive Predictive Value | 66.67% | 44.40% to 83.36% |
| Negative Predictive Value | 87.50% | 66.42% to 96.12% |
| Accuracy | 78.57% | 59.05% to 91.70% |

B Osteosarcoma vs All
| Test | Yes | n | No | n | Total |
| --- | --- | --- | --- | --- | --- |
| Yes | True Positive | 8 | False Positive | 4 | 12 |
| No | False Negative | 2 | True Negative | 42 | 44 |
| Total |  | 10 |  | 46 |  |

**Table 7. Performance of the EpiSwitch array-based biomarker classifier for calling presence of osteosarcoma in the validation cohort.**
| Results Statistics | Value | 95%CI |
| --- | --- | --- |
| Sensitivity | 80.00% | 44.39% to 97.48% |
| Specificity | 91.30% | 79.21% to 97.58% |
| Positive Likelihood Ratio | 9.2 | 3.43 to 24.67 |
| Negative Likelihood Ratio | 0.22 | 0.06 to 0.76 |
| Disease prevalence | 17.86% | 8.91% to 30.40% |
| Positive Predictive Value | 66.67% | 42.72% to 84.28% |
| Negative Predictive Value | 95.45% | 85.84% to 98.64% |
| Accuracy | 89.29% | 78.12% to 95.97% |

The multi-choice calls for canine malignant melanoma demonstrated high sensitivity of 77.78%, with no false positive calls and accuracy against healthy controls at 92.6% and accuracy against all the indications included in the validation cohort at 96.4% (Table 8). Top markers used in the canine malignant melanoma individual classifier are listed in the Supplemental Table 2.

**Table 8.**
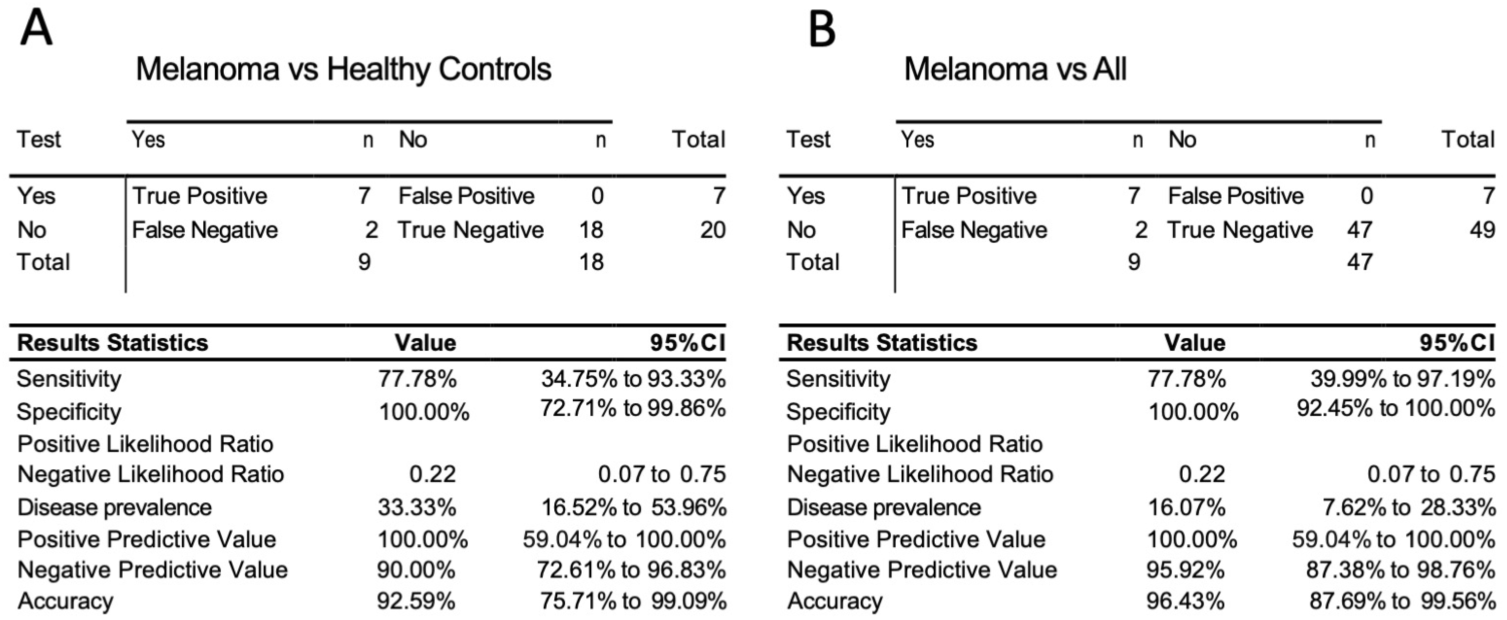
Performance of the EpiSwitch array-based biomarker classifier for calling presence of canine malignant melanoma in the validation cohort. Confusion matrix and test performance statistics for the multi-choice classifier against just healthy controls (A) and the full cohort of 56 samples, including 2 lymphomas, threes sarcomas and melanoma.

### Case Study

One of the first case studies for the developed multi-step classifier concerned a Golden Retriever SCB140842.

In February 2020, at the age of 4 years and 3 months SCB140842 was diagnosed with intranasal, poorly differentiated malignant neoplasm with vascular invasion. The nasal biopsy results indicated a very aggressive tumor as noted by the poor level of differentiation, moderate to high cancer cell activity (mitotic index), and cancer cell invasion into the blood vessels. A first blood sample CANIS080 was collected and processed on *EpiSwitch* array platform at this point.

As a treatment SCB140842 underwent Stereotactic radiation therapy (SRT) + chemotherapy (Doxorubicin), with reported 1-1.5 years of survival on average, with a moderate risk of intermittent to chronic rhinitis.

SCB140842 has developed rhinitis linked to the inflammation and remodeling that has occurring at the site of radiation. He was given Prednisone and Clindamycin for the side effects. Following the full cycle of treatment, a second blood sample CANIS193 was collected and processed on *EpiSwitch* array platform at that point.

In October 2023, with no symptoms of any complications, SCB140842 has undergone a recheck CT scan which confirmed no evidence of regrowth of his right sided nasal mass, and no evidence of spread to his lungs.

At the same time CT scan revealed a caudodorsal mediastinal mass, and a retroperitoneal mass. The mass within his chest was displacing the aorta and compressing the azygous vein. The mass within his abdomen was surrounding the cranial mesenteric artery. Both masses were located close to blood vessels, so performing a fine needle aspirate with cytology would have been too risky and dangerous. A third blood sample CANISOJB was collected and processed at this point.

Applying the multi-choice EpiSwitch classifier test, described earlier, to the longitudinal set of three samples, the results were the following:

1. First sample collection. A strong call for cancer, but only by the lymphoma general classifier for the first sample CANIS080 collected after the initial diagnosis: Lymphoma as a class - 0.72103471; Control - 0.27896529, Cut-off >=0.6. None of the two individual lymphomas showed a strong match. Independent clinical diagnosis: poorly differentiated malignant tumour.
2. Second sample collection. Improved call for healthy state by both lymphoma and sarcoma general classifiers for the second sample CANIS193, collected after the completion of the initial treatment. Independent clinical diagnosis: no evidence of regrowth of nasal mass.
3. Third sample collection. A strong call for cancer by the sarcoma general classifier for the third sample CANISOJB, collected after the second diagnosis in October 2023: sarcoma as a class - 0.72687131, Control - 0.27312869, Cut-off >=0.6;

Following up with the individual indication classifiers for CANISOJB, produced a strongest call for HSA across all indications - 0.83977288.

Independent clinical diagnosis: Newly identified caudodorsal mediastinal mass in conjunction with a retroperitoneal mass outlining the cranial mesenteric/celiac arteries. performing a fine needle aspirate with cytology was too risky and dangerous. Findings may represent atypical metastatic disease from the historical nasal anaplastic sarcoma.

### Network regulation analysis

The *EpiSwitch* profiling, apart from delivering robust biomarker modality, also provides invaluable insight into high level integrated network regulation, as reflected in systemic readouts^48,62^. Similarly, to the analysis of the previous applications in human biology, identified network of 3D genomic canine biomarkers is directly linked to the genomic loci they modulate, providing insights into affected genes and pathways.

For example, pathway enrichment for the closest coding regions for the top *EpiSwitch* sarcoma markers, common across HSA, histiocytic sarcoma, and osteosarcoma, identified a number of affected Super Pathways, as listed in Fig. 4.

**Figure 4.**
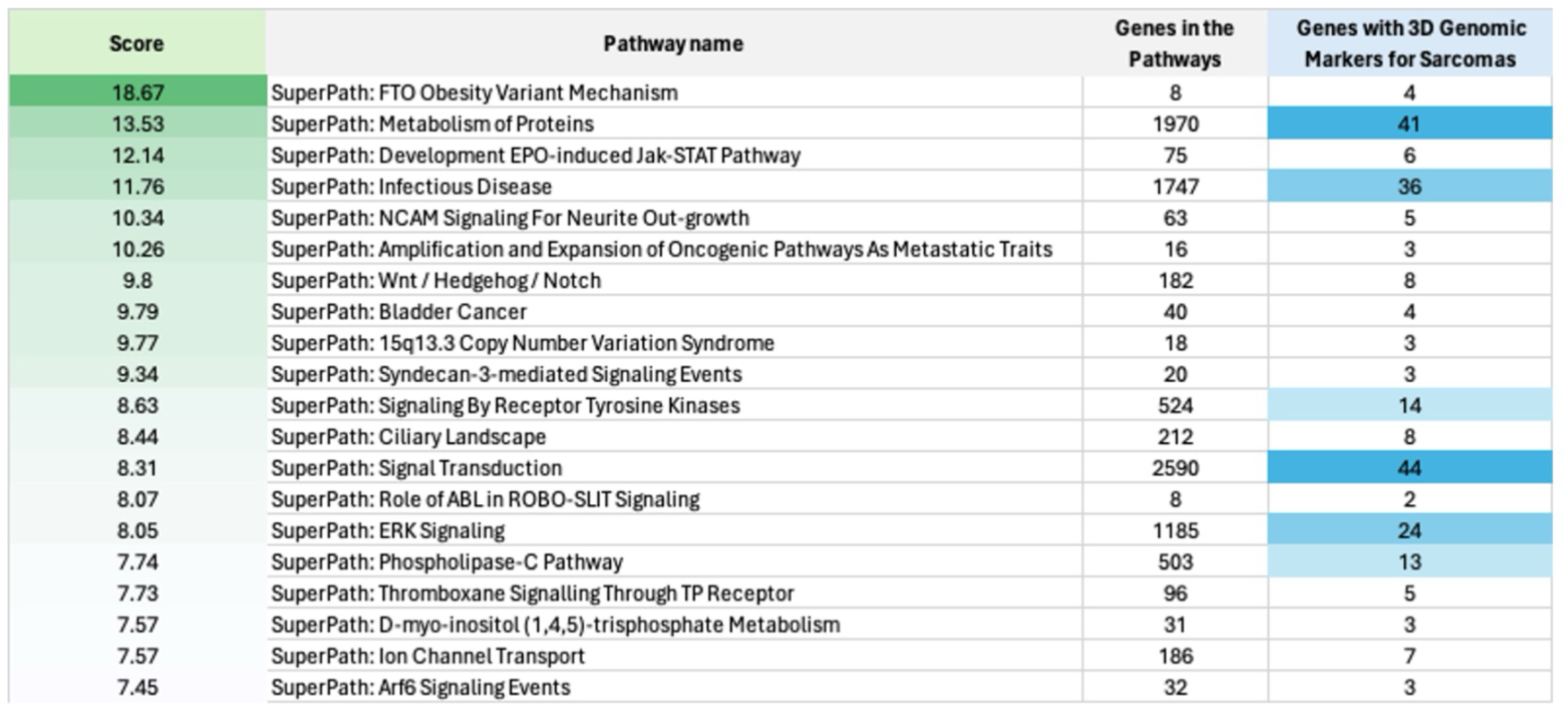
Mapping of the top 100 systemic 3D genomic common sarcoma markers to biological pathways. Analysis of the top 3D genomic markers common between HSA, histiocytic sarcoma, and osteosarcoma.

Among those, the Super Pathway for ERK Signalling, reveals 24 genes modulated by common *EpiSwitch* 3D genomic architecture. This Super Pathway also includes the pathway for Molecular Mechanisms of Cancer, with 19 genes modulated by the *EpiSwitch* 3D sarcoma markers. Those include ARGEF7, CD4, GAB1, CCN2, HAPLN1, GNB1L, NRG3, COL19A1, COL4A1, COL4A2, WNT5A, BMP1, SRC, CXCR2, CXCR4, VEGFC, VHL, VCAN, FLT3. Interestingly, the role of SRC in solid tumours and the importance of Src signalling in sarcomas are well documented, as well as recent insights into the WNT role in soft tissue sarcomas^73–75^.

Furthermore, we have also analysed the footprints of 3D genomic networks for each of the canine cancer indications. With such data over-imposed against the gene map, we have followed it up to a by building STRING protein interaction networks^72^ showing interactions at the protein that are deduced through 3D genomic architecture, all captured at the systemic level.

The STRING protein network for canine DLBCL, for example, shows a key group of affected proteins, such as MYC, BDNF, LRRK2, NRXN1, HDAC2. (Figure 5), all consistent with the human DLBCL cases^76–78^.

**Figure 5.**
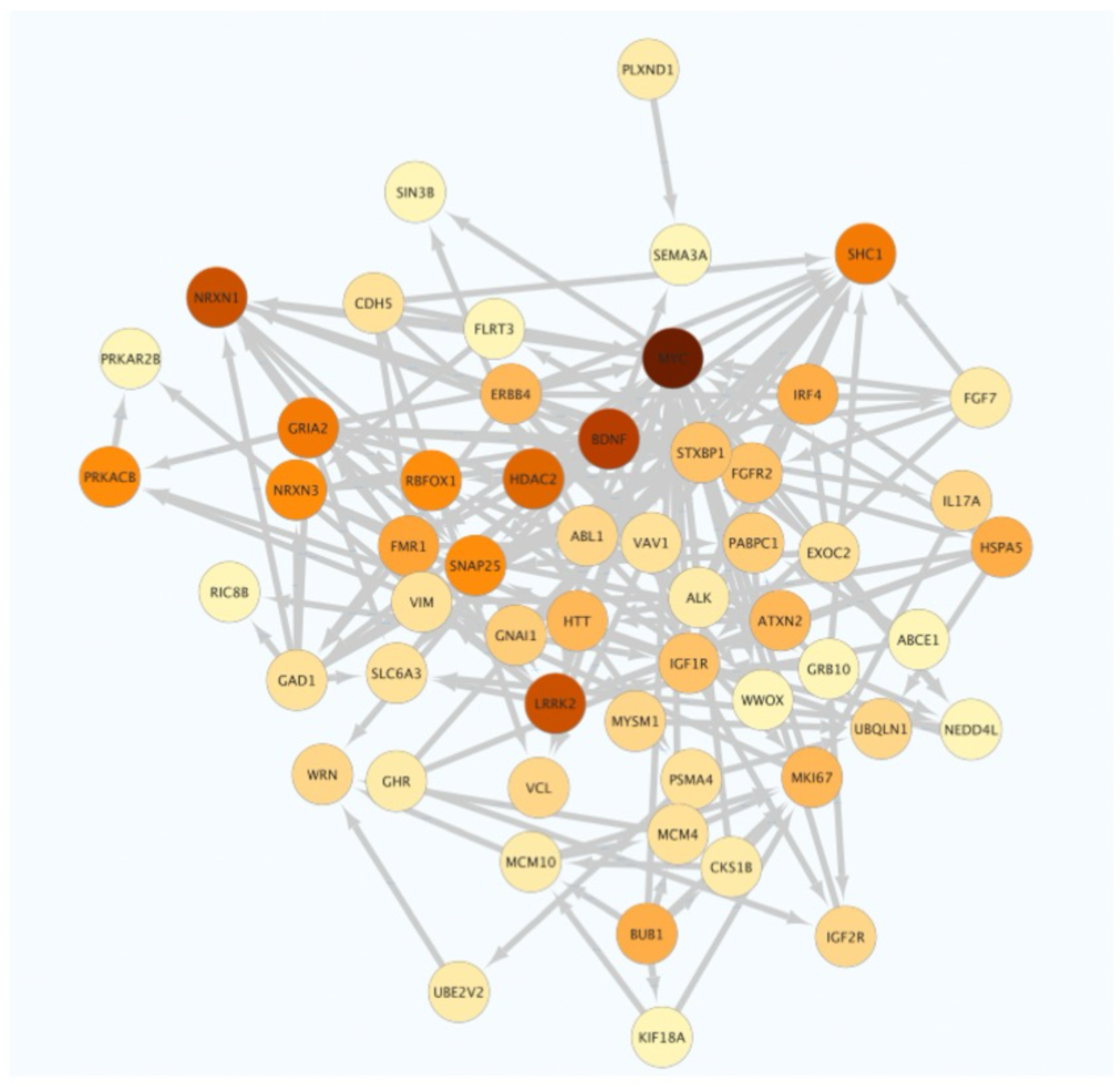
String network of DLBCL systemic profile, captured through the top 200 EpiSwitch markers. String network for DLBCL shows >550 affected gene products, with the nodes shown for over 10 connections. The density of colour corresponds to the number of connections, leading with MYC, BDNF, LRRK2, NRXN1, HDAC2.

The protein network for canine TZL, for example, shows a key group of modulated proteins, such as *ESR1, GATA4, CD44, CRBP1, ACTA1* (Figure 6), all consistent with the human T-cell NHL cases^79–81^.

**Figure 6.**
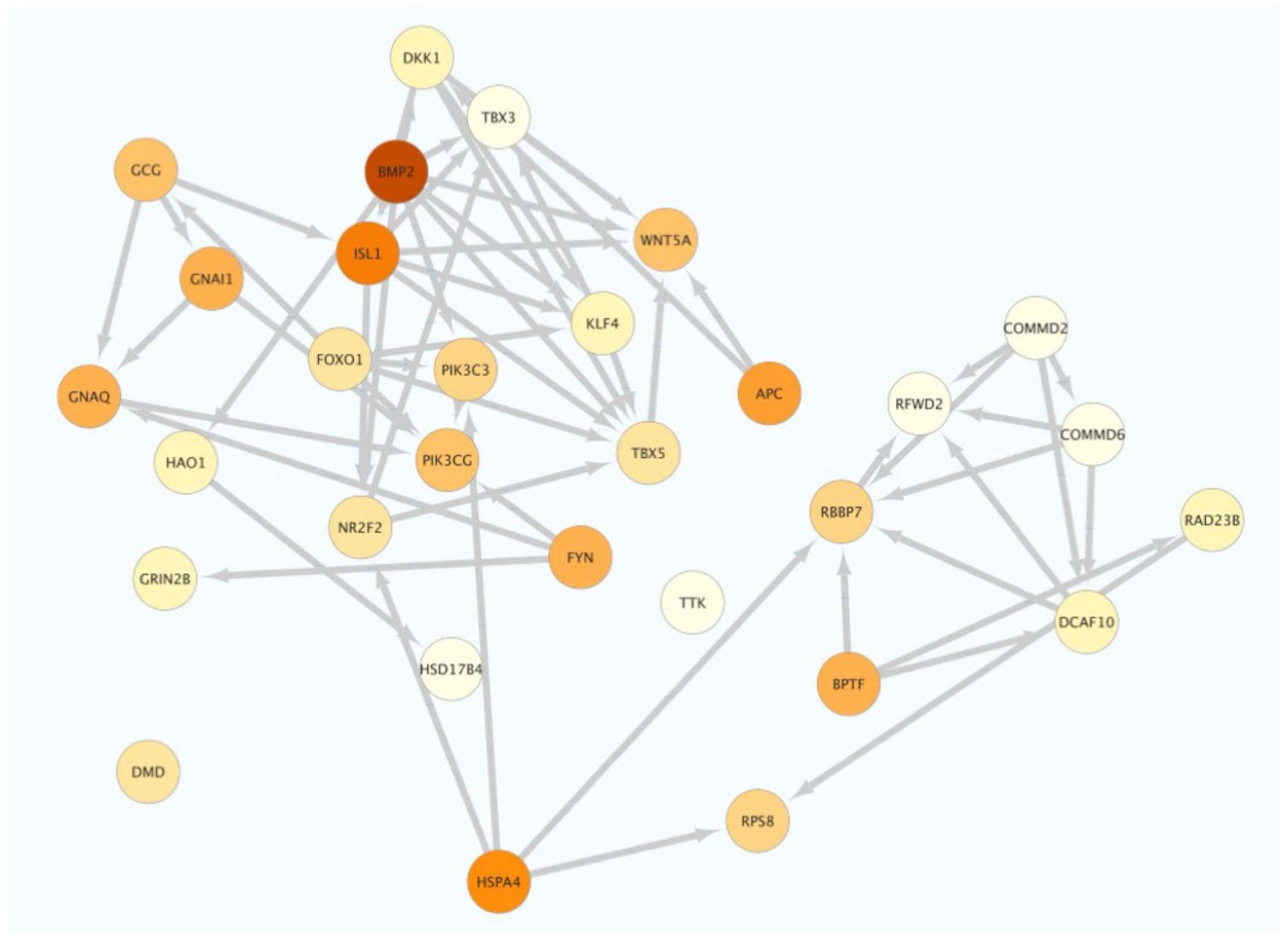
String network of TZL systemic profile, captured through the top 200 EpiSwitch markers. String network for TZL shows affected gene products, with the nodes shown for over 10 connections. The density of colour corresponds to the number of connections, leading with BMP2, ISL1, APC, HSPA4, FYN, BPTF.

The protein network for canine HSA, for example, shows a key group of modulated proteins, such as MYC, EGFR, POLR2B, PTPRD, NTRK2, RUNX2 (Figure 7), all consistent with the human angiosarcoma cases^82–84^.

**Figure 7.**
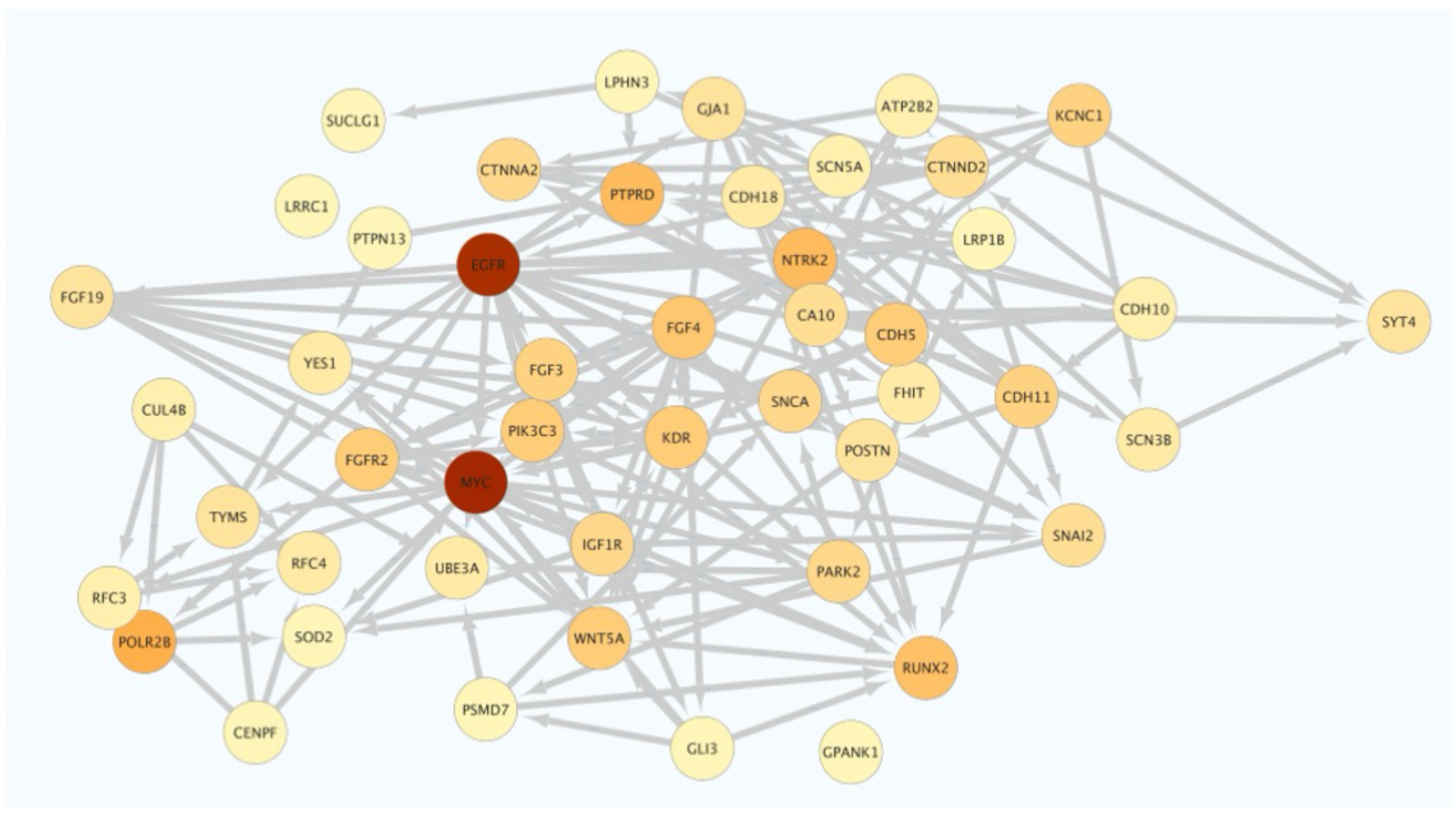
String network of HSA systemic profile, captured through the top 200 EpiSwitch markers. String network for HSA shows affected gene products, with the nodes shown for over 10 connections. The density of colour corresponds to the number of connections, leading with MYC, EGFR, POLR2B, PTPRD, NTRK2, RUNX2.

The protein network for canine histiocytic sarcoma, for example, shows a key group of modulated proteins, such as EZH2, EPRS, HIST1H4F, CDC6, TOP2A, CUL1, PABPC1, CDH2, NTRK2. (Figure 8), all consistent with the human histiocytic sarcoma cases^85,86^.

**Figure 8.**
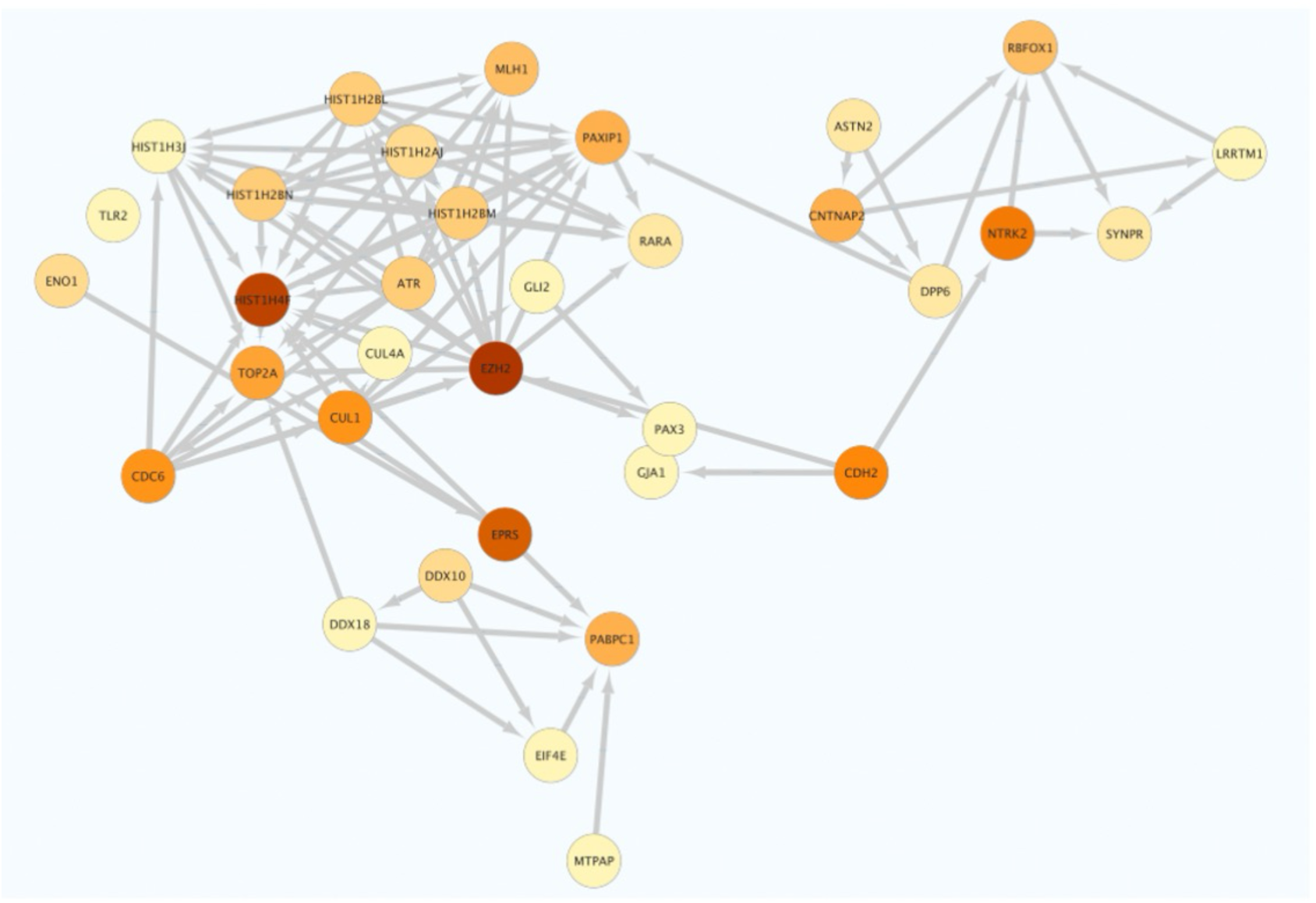
String network of histiocytic sarcoma systemic profile, captured through the top 200 EpiSwitch markers. String network for histiocytic sarcoma shows affected gene products, with the nodes shown for over 10 connections. The density of colour corresponds to the number of connections, leading with EZH2, EPRS, HIST1H4F, CDC6, TOP2A, CUL1, PABPC1, CDH2, NTRK2.

The protein network for canine osteosarcoma, for example, shows a key group of modulated proteins, such as EGFR, IL17A, CA10, WASL, SH3GL2, POLR2B (Figure 9), all consistent with the human osteosarcoma cases^87–89^.

**Figure 9.**
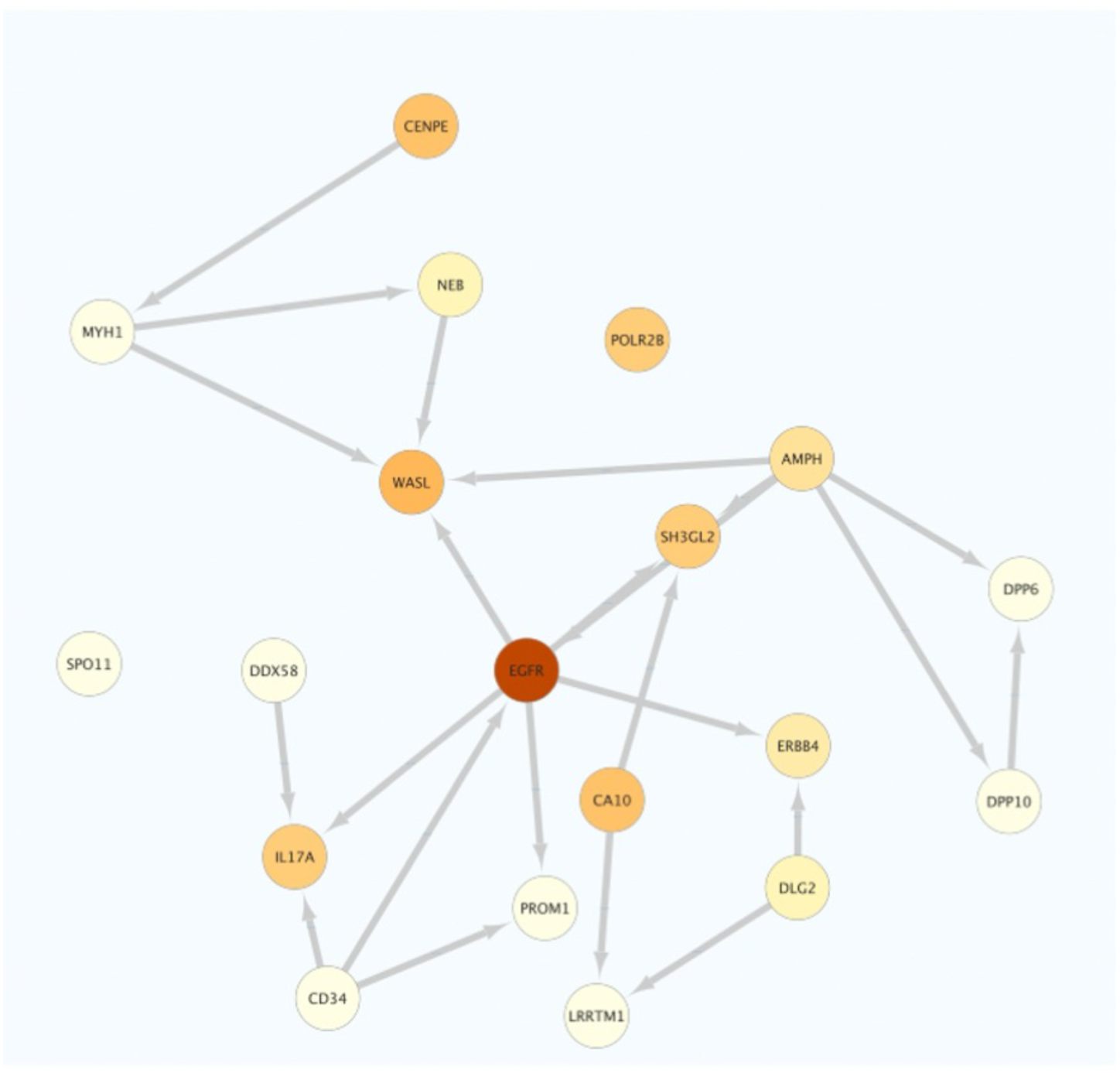
String network of osteosarcoma systemic profile, captured through the top 200 EpiSwitch markers. String network for osteosarcoma shows affected gene products, with the nodes shown for over 10 connections. The density of colour corresponds to the number of connections, leading with EGFR, IL17A, CA10, WASL, SH3GL2, POLR2B.

The protein network for canine malignant melanoma, for example, shows a key group of modulated proteins, such as ESR1, GATA4, CD44, CTBP1, ACTA1 (Figure 10) all consistent with the human and canine malignant melanoma cases^90–92^.

**Figure 10.**
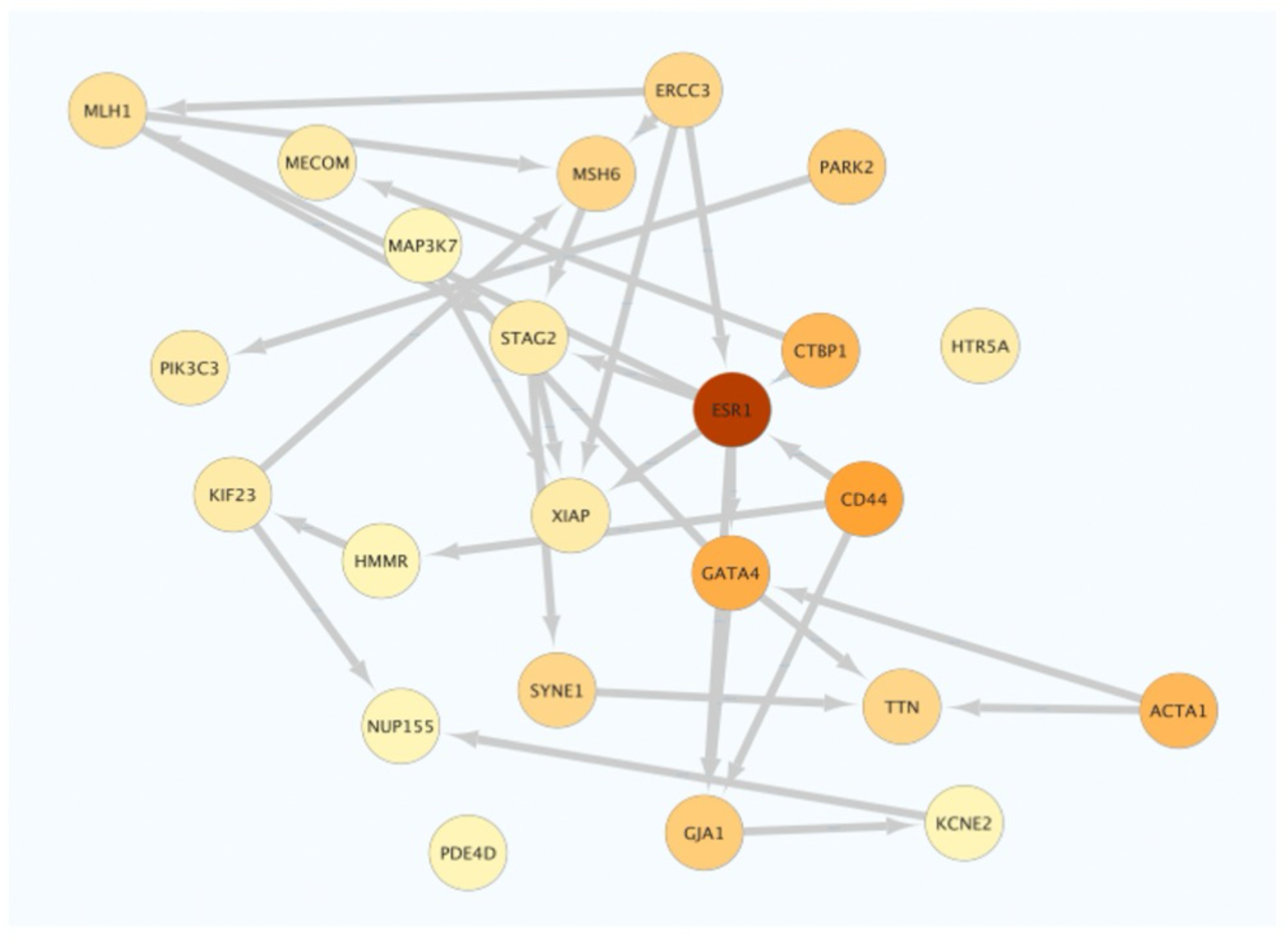
String network of canine malignant melanoma systemic profile, captured through the top 200 EpiSwitch markers. String network for canine malignant melanoma shows >550 affected gene products, with the nodes shown for over 10 connections. The density of colour corresponds to the number of connections, leading with ESR1, GATA4, CD44, CTBP1, ACTA1.

It is important to emphasize, that any observed redundancy at the level of protein network interaction is a reductional representation of distinct and unique conditional chromosome conformations within the complex 3D architectural landscape across the same protein encoding gene locus. This reflect different contribution for 3D genomic modulation into the same gene loci regulation when different oncological indications are compared.

## Discussion

Here we describe first canine liquid biopsy application with the approach that has demonstrated earlier the strong advantages of systemic 3D genomic profiling using whole-genome *EpiSwitch* arrays in human oncological applications for diagnosis, prognosis, and prediction of response to treatment^48–51,65^. With the challenging task for a multi-choice stratification, we have built an array-based two step classifier. Extensive analysis of the systemic signatures in canine cancers revealed strong signatures by class, with markers shared by all tested lymphomas for one class and distinct strong markers shared by all tested sarcomas for another class. In light of this, we are of opinion, that at the level of 3D genomics the systemic signatures, with all its integration of genetic and epigenetic inputs^52^, systemic profiling of cancers does not reveal itself as a universal pan-cancer signature, but as a distinct family of signatures by class, reflecting their distinct mechanisms, shared, in our example, by sarcomas.

Thus, firstly, in the context of the multi-choice classifier, the profile of an individual sample undergoes class stratifications against heathy controls. The second stage of classification benefits from the individual stratification models, based on unique selected markers for each individual indication.

Today, this approach has been tested on 56 independent samples, representing a mixture of healthy controls, two subtypes of lymphoma, three different types of sarcoma and canine malignant melanoma. The results of the first validation demonstrate high efficacy of stratification. Importantly, they also demonstrate high specificity against all other cancer indications. This is an important advantage against genetic mutation approaches in free cancer circulating DNA, which shares similar mutational profile between many cancers. For example, PetDx OncoK9 multi-cancer early detection test for detection of cancer associated genomic alterations in DNA isolated from canine blood using next generation sequencing is able to detect genetic alterations common for 30 cancers with high accuracy of sequencing readout, but according to the ONCOK9 Test Interpretation Guide the report outcome is specified in three types: 1) cancer signal detected, cancer signal not detected, and not reportable – sample failed ^93^. Example of the SCB140842 case study has demonstrated highly informative longitudinal changes in specificity captured by the *EpiSwitch* profiling after the radiotherapy and chemotherapy treatments, as well as after the second diagnosis with unavailable pathology results.

Current results constitute the first systemic oncological classifications as an *EpiSwitch* specific canine blood-based (EpiSwitch SCB) test. We anticipate to significantly expand the list of the individual indications covered by the test and to extend the validation exercise to all underrepresented cases. Using the modular structure for the test, we aim to add individual classifiers to the existing profiles, at the level of data analysis, without any changes to the whole-genome EpiSwitch canine array design. Given the feedback form the veterinary practices, we strongly believe that the developed *EpiSwitch* SCB™ test approach will be of interest for veterinary specialist as an additional tool contributing to the informed decisions on diagnosis, treatment, and best standards of care for our canine companions.

## Conclusions

Clearly, there is a pressing need to develop better non-invasive (blood) biomarker assays to assess early canine oncological indications in advance of therapeutic intervention. Here we report on a novel 3D genomics approach to identify systemic blood-based markers for canine DLBCL, TZL, HSA, histiocytic sarcoma, osteosarcoma, and canine malignant melanoma in an assay format that encompasses multiple classes and phenotypes of cancer. The approach described here is based on earlier applications in human oncology. As a non-invasive, blood-based test, *EpiSwitch* SCB, promises to assist veterinary specialists in diagnosis of disease and associated treatment decisions, to better utilize alternative effective treatments, minimize or avoid unnecessarily toxicity, and efficiently manage costs and resources.

## Authors contributions

EH, AA conceived the study, MS assisted with study design and clinical samples collection. AD, MS, EH assisted with *EpiSwitch* array design. MS and RP assisted with design of experiments. JG and AA planned and reviewed experiments. TN, DV, KS, EH analysed the data. AD, MI, CW, AH, AG performed experiments. SF helped with interpretation of the data and writing of the manuscript. JFM obtained the samples. AA, EH, AB, TG and JFM wrote and reviewed the manuscript.

## Acknowledgements

The authors would like to thank past and present members of OBD for their operational support, Mitzi Lewellen in managing samples transfers, all inventory activities, and metadata collection for the dogs in the study, Olly Hunter for his assistance in data analysis. In addition, we acknowledge Agilent Technologies, Inc. for supply of *EpiSwitch* designed CGH microarray slides.

The authors want to express their gratitude to the SCB140842 family for their permission to share the case study details.

