## Supplemental Table 1 for "Whole Genome 3D Blood Biopsy Profiling of Canine Cancers: Development and Validation of EpiSwitch Multi-Choice Array-Based Diagnostic Test"

| **Sample ID** | Phenotype | Age at Diagnosis Sample | Sex Neuter Status | Breed | Breed Median LifeSpan |
|---|---|---|---|---|---|
| CANIS092 | DLBCL | 5.9 | MN | Hound Mix | n/a |
| CANIS107 | DLBCL | 7.7 | MN | Beagle | 10.9 |
| CANIS086 | DLBCL | 12.6 | M? | West Highland white terrier | 13.8 |
| CANIS109 | DLBCL | 13 | FS | Labrador retriever | 12.5 |
| CANIS096 | DLBCL | 9.6 | MN | Golden Retriever | 12.5 |
| CANIS108 | DLBCL | 7.0 | FS | Bullmastiff | 8.5 |
| CANIS106 | DLBCL | 11.6 | FS | German Shepherd Dog | 12.6 |
| CANIS090 | DLBCL | 9.8 | FI | Standard Poodle | 11.96 |
| CANIS104 | DLBCL | 4.7 | MN | Vizsla | 11 |
| CANIS097 | DLBCL | 3.8 | MN | German Shepherd Dog | 12.6 |
| CANIS095 | DLBCL | 16.1 | FS | Border Collie/Sheltie Hix | 13.7 |
| CANIS091 | DLBCL | 11.7 | MN | Labrador Retriever | 12.5 |
| CANIS094 | DLBCL | 6.2 | MN | Golden Retriever | 12.5 |
| CANIS098 | DLBCL | 8.8 | MN | Labrador retriever | 12.5 |
| CANIS103 | DLBCL | 11.5 | FS | Springer Spaniel Hix | 10.7 |
| CANIS085 | DLBCL | 14.0 | MN | Golden Retriever | 12.5 |
| CANIS087 | DLBCL | 8.8 | MI | Golden Retriever | 12.5 |
| CANIS099 | DLBCL | 5.7 | FS | Labradoodle | 8.5 |
| CANIS102 | DLBCL | 3.8 | MN | Standard Poodle | 11.96 |
| CANIS093 | DLBCL | 3.1 | MN | Mastiff | 9.053958476 |
| CANIS257 | Hemangiosarcoma | 12.7 | FS | Portuguese Water Dog | 6.577629476 |
| CANIS276 | Hemangiosarcoma | 10.3 | FS | Golden retriever | 12.51186402 |
| CANIS285 | Hemangiosarcoma | 11.2 | FS | Golden retriever | 12.51186402 |
| CANIS297 | Hemangiosarcoma | 11.7 | FS | German Shepherd Dog | 10.6939311 |
| CANIS266 | Hemangiosarcoma | 9.2 | FI | Golden retriever | 12.51186402 |
| CANIS273 | Hemangiosarcoma | 8.3 | FI | Golden retriever | 12.51186402 |
| CANIS309 | Hemangiosarcoma | 13 | MN | Golden retriever | 12.51186402 |
| CANIS260 | Hemangiosarcoma | 7.8 | MN | Portuguese Water Dog | 6.577629476 |
| CANIS294 | Hemangiosarcoma | 11.3 | FS | Golden retriever | 12.51186402 |
| CANIS279 | Hemangiosarcoma | 9.5 | FS | Gordon setter | 10.23397217 |
| CANIS263 | Hemangiosarcoma | 6.7 | MN | Golden retriever | 12.51186402 |
| CANIS282 | Hemangiosarcoma | 10.1 | FS | Boxer | 10.39835729 |
| CANIS303 | Hemangiosarcoma | 9.1 | MN | Golden retriever | 12.51186402 |

| | | | | | |
|---|---|---|---|---|---|
| CANIS269 | Hemangiosarcoma | 14.5 | F | Golden retriever | 12.51186402 |
| CANIS254 | Hemangiosarcoma | 10 | MN | Beagle | 10.90200776 |
| CANIS291 | Hemangiosarcoma | 8 | MN | Golden retriever | 12.51186402 |
| CANIS288 | Hemangiosarcoma | 8.4 | MN | Golden retriever | 12.51186402 |
| CANIS312 | Hemangiosarcoma | 9 | MN | Australian shepherd | 14.05612594 |
| CANIS306 | Hemangiosarcoma | 10 | MN | Mixed breed | 12.54757016 |
| CANIS300 | Hemangiosarcoma | 9.3 | FS | Golden retriever | 12.51186402 |
| CANIS178 | Histiocytic Sarcoma | 7 | MN | Golden Retriever | 12.51186402 |
| CANIS173 | Histiocytic Sarcoma | 8.0 | MI | Golden Retriever | 12.51186402 |
| CANIS167 | Histiocytic Sarcoma | 4.9 | FI | Golden Retriever | 12.51186402 |
| CANIS186 | Histiocytic Sarcoma | 12.4 | FS | Labrador Retriever | 12.32842802 |
| CANIS190 | Histiocytic Sarcoma | 11.4 | FS | German wire-haired pointer | 12.57494867 |
| CANIS179 | Histiocytic Sarcoma | 8.3 | MI | Golden Retriever | 12.51186402 |
| CANIS189 | Histiocytic Sarcoma | 10 | MN | German shepherd Hix | 11.92465207 |
| CANIS165 | Histiocytic Sarcoma | 6.0 | MI | Bernese Mountain Dog | 8 |
| CANIS187 | Histiocytic Sarcoma | 6 | FI | Golden Retriever | 12.51186402 |
| CANIS180 | Histiocytic Sarcoma | 7.4 | FS | MIX: BMD x Australian shepherd | 7.787759526 |
| CANIS176 | Histiocytic Sarcoma | 9.8 | MI | Bernese Mountain Dog | 8 |
| CANIS171 | Histiocytic Sarcoma | 6.5 | MN | Rottweiler | 9.327857632 |
| CANIS160 | Melanoma | 13.6 | MN | Labrador Retriever | 12.5 |
| CANIS125 | Melanoma | 13.1 | M? | Golden Retriever | 12.5 |
| CANIS188 | Melanoma | 4 | FI | briard | 12 |
| CANIS166 | Melanoma | 8.3 | MN | Keeshond/Chow Mix | 13 |
| CANIS128 | Melanoma | 11.9 | F? | Miniature Schnauzer | 11 |
| CANIS174 | Melanoma | 11.9 | FS | Golden Retriever | 12.5 |
| CANIS147 | Melanoma | 14.0 | M? | Golden Retriever | 12.5 |
| CANIS142 | Melanoma | 5.0 | F? | Doberman Pinscher | 5.3 |
| CANIS136 | Melanoma | 13.7 | FS | Rhodesian Ridgeback | 11 |
| CANIS141 | Melanoma | 11.7 | F? | Shetland Sheepdog | 12.4 |
| CANIS149 | Melanoma | 12.1 | MN | Golden Retriever | 12.5 |
| CANIS154 | Melanoma | 9.2 | MI | Golden Retriever | 12.5 |
| CANIS161 | Melanoma | 13.0 | FS | German Shorthair Pointer | ?? |
| CANIS159 | Melanoma | 13.2 | FS | Mix | 12.7 |
| CANIS143 | Melanoma | 11.0 | M? | Miniature Schnauzer | 11 |

| | | | | | |
|---|---|---|---|---|---|
| CANIS150 | Melanoma | 8.8 | F? | Yorkshire terrier | 12.77 |
| CANIS138 | Melanoma | 12.9 | MN | Labrador Retriever | 12.5 |
| CANIS129 | Melanoma | 4.0 | FS | Basenji mix | ?? |
| CANIS177 | Melanoma | 2.2 | FI | Doberman pinscher | 5.3 |
| CANIS214 | Osteosarcoma | 0.4 | MN | Rhodesian ridgeback | 11.10746064 |
| CANIS238 | Osteosarcoma | 3.5 | FS | Labrador retriever | 12.32842802 |
| CANIS232 | Osteosarcoma | 5.2 | FS | Mastiff | 9.053958476 |
| CANIS245 | Osteosarcoma | 12.2 | MN | Golden retriever | 12.51186402 |
| CANIS223 | Osteosarcoma | 7.7 | FS | Newfoundland | 9.590691308 |
| CANIS229 | Osteosarcoma | 5.9 | MN | Golden retriever | 12.51186402 |
| CANIS248 | Osteosarcoma | 8.4 | MN | Akita | 8.640543007 |
| CANIS220 | Osteosarcoma | 6.3 | FS | Rottweiler | 9.327857632 |
| CANIS241 | Osteosarcoma | 1.9 | FI | Rhodesian ridgeback | 11.10746064 |
| CANIS235 | Osteosarcoma | 7 | MN | Rottweiler | 9.327857632 |
| CANIS251 | Osteosarcoma | 11 | MN | Mixed breed | 12.54757016 |
| CANIS202 | Osteosarcoma | 13.1 | FS | Irish setter | 11.59479809 |
| CANIS196 | Osteosarcoma | n/a | F | Rottweiler | 9.327857632 |
| CANIS217 | Osteosarcoma | 3.9 | F | Rottweiler | 9.327857632 |
| CANIS205 | Osteosarcoma | 5.9 | FS | Saint Bernard | 7.753479352 |
| CANIS211 | Osteosarcoma | 11.5 | FS | Golden retriever | 12.51186402 |
| CANIS243 | Osteosarcoma | 7.6 | FS | Rottweiler | 9.327857632 |
| CANIS208 | Osteosarcoma | 8.5 | FS | Great Dane | 7.978097194 |
| CANIS199 | Osteosarcoma | 3 | FI | Golden retriever | 12.51186402 |
| CANIS226 | Osteosarcoma | 8.4 | FS | Labrador retriever | 12.32842802 |
| CANIS182 | TZL | 9.3 | MN | Golden Retriever | 12.51186402 |
| CANIS168 | TZL | 8.1 | FS | Labrador/Boxer Mix | 13.449692 |
| CANIS140 | TZL | 9.0 | M? | Golden Retriever | 12.51186402 |
| CANIS164 | TZL | 11.9 | FS | Golden Retriever | 12.51186402 |
| CANIS158 | TZL | 8.9 | FS | Golden Retriever | 12.51186402 |
| CANIS172 | TZL | 5.5 | MN | Bernese Mountain Dog | 8 |
| CANIS132 | TZL | 9.9 | FS | Golden Retriever | 12.51186402 |
| CANIS139 | TZL | 6.7 | MN | Golden Retriever | 12.51186402 |
| CANIS185 | TZL | 5.4 | FS | Portuguese Water Dog | 6.577629476 |
| CANIS153 | TZL | 9.0 | MN | Golden Retriever | 12.51186402 |

| | | | | | |
|---|---|---|---|---|---|
| CANIS145 | TZL | 8.0 | MN | Mastiff | 9.053958476 |
| CANIS123 | TZL | 8.7 | FS | Golden Retriever | 12.51186402 |
| CANIS127 | TZL | 11.2 | MI | Golden Retriever | 12.51186402 |
| CANIS137 | TZL | 7.5 | M? | Mastiff | 9.053958476 |
| CANIS146 | TZL | 9.6 | F? | Golden Retriever | 12.51186402 |
| CANIS124 | TZL | 10.4 | M? | Golden Retriever | 12.51186402 |
| CANIS122 | TZL | 9.1 | M? | Golden Retriever | 12.51186402 |
| Canis0064 | Control | n/a | n/a | n/a | n/a |
| Canis0048 | Control | n/a | n/a | n/a | n/a |
| Canis0058 | Control | n/a | n/a | n/a | n/a |
| Canis0060 | Control | n/a | n/a | n/a | n/a |
| Canis0065 | Control | n/a | n/a | n/a | n/a |
| Canis0076 | Control | n/a | n/a | n/a | n/a |
| Canis0043 | Control | n/a | n/a | n/a | n/a |
| Canis0041 | Control | n/a | n/a | n/a | n/a |
| Canis0057 | Control | n/a | n/a | n/a | n/a |
| Canis0068 | Control | n/a | n/a | n/a | n/a |
| Canis0051 | Control | n/a | n/a | n/a | n/a |
| Canis0074 | Control | n/a | n/a | n/a | n/a |
| Canis0070 | Control | n/a | n/a | n/a | n/a |
| Canis0045 | Control | n/a | n/a | n/a | n/a |
| Canis0072 | Control | n/a | n/a | n/a | n/a |
| Canis0054 | Control | n/a | n/a | n/a | n/a |
| Canis0079 | Control | n/a | n/a | n/a | n/a |
| Canis0038 | Control | n/a | n/a | n/a | n/a |
| CANIS093 | Control | n/a | n/a | n/a | n/a |
| CANIS324 | Control | n/a | M | Golden retriever | 12.51186402 |
| CANIS321 | Control | 11.6 | FS | Golden retriever | 12.51186402 |
| CANIS333 | Control | n/a | F | Golden retriever | 12.51186402 |
| CANIS336 | Control | 4 | M | Golden retriever | 12.51186402 |
| CANIS345 | Control | 10.3 | FS | Golden retriever | 12.51186402 |
| CANIS315 | Control | 5.2 | MN | Golden retriever | 12.51186402 |
| CANIS373 | Control | 15.5 | MN | German Shepherd Dog | 10.6939311 |
| CANIS364 | Control | 4 | F | Rottweiler | 9.327857632 |

| | | | | | |
|---|---|---|---|---|---|
| CANIS361 | Control | 5.3 | FI | Doberman pinscher | 9.371606206 |
| CANIS367 | Control | 6.6 | F | Great Pyrenees | 10.06291353 |
| CANIS342 | Control | 7.6 | FS | Golden retriever | 12.51186402 |
| CANIS351 | Control | 5.2 | M | Rottweiler | 9.327857632 |
| CANIS318 | Control | n/a | n/a | Great Pyrenees | 10.06291353 |
| CANIS348 | Control | n/a | n/a | Rottweiler | 9.327857632 |
| CANIS370 | Control | 3.9 | M | Rottweiler | 9.327857632 |
| CANIS354 | Control | 6.9 | FS | Chesapeake Bay Retriever | 8.057437829 |
| CANIS330 | Control | 5.5 | FS | Golden retriever | 12.51186402 |
| CANIS339 | Control | 5.5 | n/a | Golden retriever | 12.51186402 |
| CANIS358 | Control | 9.1 | FI | Golden retriever | 12.51186402 |
| CANIS327 | Control | 3.6 | F | Great Pyrenees | 10.06291353 |
| CANIS080 | Case Study_1st Sample | 4.3 | M | Golden Retriever | 12.51186402 |
| CANIS193 | Case Study_2nd Sample | 5.5 | M | Golden Retriever | 12.51186402 |
| CANIS381 | Case Study_3nd Sample | 7 | M | Golden Retriever | 12.51186402 |
