## Supplemental Table 2 for "Whole Genome 3D Blood Biopsy Profiling of Canine Cancers: Development and Validation of EpiSwitch Multi-Choice Array-Based Diagnostic Test"

**Top Markers for Lymphomas as a Class**

| 3D array probe | logFC | Average Expression | Moderated t-test for LIMMA | P Value | Adjusted P Value | B-statistics (log-odds) for LIMMA | Fold Change (FC) |
|---|---|---|---|---|---|---|---|
| CanFam3_5_11838594_11840423_11964855_11970127_RR | -0.333282041 | 8.674710737 | -3.333108105 | 0.00180394 | 0.836524557 | -1.364057356 | -1.259876257 |
| CanFam3_21_28553546_28564169_28793102_28796106_RR | -0.290273118 | 10.29873137 | -3.332741598 | 0.00180582 | 0.836524557 | -1.364850973 | -1.222871759 |
| CanFam3_14_43698959_43704181_43837511_43840004_FF | -0.301224793 | 8.258254003 | -3.275767825 | 0.00212083 | 0.836524557 | -1.487684241 | -1.232190052 |
| CanFam3_38_13144583_13147670_13283324_13285698_RF | -0.285259109 | 8.465155224 | -3.14234837 | 0.00307483 | 0.836524557 | -1.771001166 | -1.218629111 |
| CanFam3_21_28553546_28564169_28705922_28713313_RR | -0.273186714 | 10.28350149 | -3.075359146 | 0.00369487 | 0.836524557 | -1.910842122 | -1.208474235 |
| CanFam3_18_20581111_20590377_20824245_20832056_RR | -0.285989552 | 8.284126668 | -2.954423505 | 0.0051223 | 0.836524557 | -2.158953796 | -1.219246264 |
| CanFam3_31_12192736_12202811_12255709_12263471_FF | 0.27578053 | 8.117683978 | 2.95345596 | 0.00513557 | 0.836524557 | -2.160915467 | 1.2106489 |
| CanFam3_21_28553546_28564169_28775044_28776391_RR | -0.271090485 | 10.18074234 | -2.844461745 | 0.00685399 | 0.836524557 | -2.379425362 | -1.206719602 |
| CanFam3_18_19956797_19962368_20072595_20076616_FR | 0.263784608 | 9.792168687 | 2.774017964 | 0.00823492 | 0.836524557 | -2.517958522 | 1.200624163 |
| CanFam3_10_51757418_51762484_51967024_51972491_RF | 0.314886168 | 8.228060541 | 2.700825628 | 0.00993955 | 0.836524557 | -2.659561002 | 1.243913502 |
| CanFam3_22_25061511_25065331_25286016_25288927_FR | -0.268337355 | 8.737941457 | -2.521511189 | 0.01557877 | 0.836524557 | -2.995875164 | -1.204418986 |
| CanFam3_11_8046746_8065995_8207494_8215964_FR | 0.2881196 | 9.940240676 | 2.51897692 | 0.01567613 | 0.836524557 | -3.000516118 | 1.221047734 |
| CanFam3_6_69242157_69243830_69293503_69294903_FR | 0.267404764 | 10.9127046 | 2.514842998 | 0.01583614 | 0.836524557 | -3.008079582 | 1.203640674 |
| CanFam3_6_57622297_57626700_57776562_57786625_FR | -0.312718308 | 8.766717897 | -2.486293331 | 0.01698244 | 0.836524557 | -3.060079538 | -1.242045744 |
| CanFam3_19_52765842_52768809_52778874_52780239_RF | -0.33582226 | 8.831207997 | -2.472529878 | 0.01756157 | 0.836524557 | -3.08500056 | -1.262096532 |
| CanFam3_18_20581111_20590377_20621045_20633611_RR | -0.264040315 | 8.382496445 | -2.469529095 | 0.01769019 | 0.836524557 | -3.090421149 | -1.200836984 |
| CanFam3_1_31826881_31839714_32016857_32023684_FR | 0.288262813 | 9.283690405 | 2.462249112 | 0.0180058 | 0.836524557 | -3.103552499 | 1.22116895 |
| CanFam3_36_2682617_2690121_2795213_2796237_RR | -0.284640851 | 8.59987041 | -2.460540187 | 0.01808062 | 0.836524557 | -3.106631053 | -1.218106987 |
| CanFam3_31_18338790_18346910_18461801_18472074_FF | 0.263269475 | 9.315444777 | 2.449106194 | 0.01858854 | 0.836524557 | -3.127190267 | 1.200195541 |
| CanFam3_22_31895855_31900418_32002824_32004507_FR | -0.393448346 | 8.230639448 | -2.432480722 | 0.01935016 | 0.836524557 | -3.156963731 | -1.313529267 |
| CanFam3_1_31913537_31919259_32016857_32023684_FR | 0.264014692 | 9.077890844 | 2.400271145 | 0.02090656 | 0.836524557 | -3.214236186 | 1.200815656 |
| CanFam3_31_33405585_33412005_33522592_33524266_RR | -0.275653561 | 9.001456194 | -2.377237726 | 0.02208794 | 0.836524557 | -3.25485768 | -1.210542358 |
| CanFam3_30_12334103_12350622_12478195_12487583_FR | 0.266849588 | 8.444742341 | 2.33613394 | 0.02434643 | 0.836524557 | -3.326645234 | 1.203177579 |
| CanFam3_36_2682617_2690121_2757079_2759175_RR | -0.277236576 | 8.519243187 | -2.222361394 | 0.03171591 | 0.836524557 | -3.520537802 | -1.211871369 |
| CanFam3_16_25979575_25996021_26150487_26154989_FF | 0.265028737 | 8.492119418 | 2.22198081 | 0.03174358 | 0.836524557 | -3.521174292 | 1.201659985 |
| CanFam3_27_25723467_25734018_25897974_25901124_RR | -0.332870588 | 8.290323553 | -2.204228043 | 0.0330582 | 0.836524557 | -3.550772868 | -1.259516994 |
| CanFam3_9_44452386_44454722_44564407_44570148_FR | 0.271247267 | 13.52100427 | 2.200690718 | 0.03332584 | 0.836524557 | -3.556649127 | 1.206850747 |
| CanFam3_X_71181031_71203919_71304170_71309996_RR | 0.266934017 | 9.364876807 | 2.197784469 | 0.03354717 | 0.836524557 | -3.561471693 | 1.203247993 |
| CanFam3_1_101457517_101462358_101690254_101692916_RR | 0.324139839 | 11.33882438 | 2.166738727 | 0.03599397 | 0.836524557 | -3.612686447 | 1.2519178 |
| CanFam3_3_25122972_25124688_25148378_25155983_FF | -0.265596971 | 8.056859049 | -2.143988296 | 0.03788579 | 0.836524557 | -3.649863797 | -1.202133376 |
| CanFam3_7_76197803_76203899_76278424_76292391_FR | -0.275249556 | 8.215072437 | -2.140472892 | 0.0381858 | 0.836524557 | -3.655581633 | -1.210203411 |
| CanFam3_25_24006585_24022607_24087831_24090714_FR | 0.268408301 | 10.3786063 | 2.129512834 | 0.03913462 | 0.836524557 | -3.673361967 | 1.204478216 |
| CanFam3_35_4962878_4965874_5028098_5034625_FR | -0.268781764 | 8.010657506 | -2.092517785 | 0.04249217 | 0.836524557 | -3.732857531 | -1.204790053 |
| CanFam3_2_78835628_78839336_78980158_78981213_FR | -0.314218976 | 8.50766289 | -2.09155622 | 0.04258271 | 0.836524557 | -3.734393142 | -1.243338371 |
| CanFam3_18_18481385_18486743_18498471_18500877_FF | -0.343861427 | 8.649918301 | -2.090901224 | 0.04264448 | 0.836524557 | -3.735438852 | -1.269148976 |
| CanFam3_1_101363857_101365401_101463964_101465898_RF | 0.310215417 | 13.01358741 | 2.087533143 | 0.04296334 | 0.836524557 | -3.740812011 | 1.239892822 |
| CanFam3_4_26771736_26778011_26987437_26991849_RF | -0.273029275 | 10.99203338 | -2.080909338 | 0.04359652 | 0.836524557 | -3.75135941 | -1.208342363 |

**Top Markers for Sarcomas as a Class**

| 3D array probe | logFC | Average Expression | Moderated t-test for LIMMA | P Value | Adjusted P Value | B-statistics (log-odds) for LIMMA | Fold Change (FC) |
|---|---|---|---|---|---|---|---|
| CanFam3_24_31264548_31267041_31327229_31330640_FR | 0.672659237 | 9.709978485 | 4.157179838 | 0.00012103 | 0.002872029 | 1.023354397 | 1.594008406 |
| CanFam3_3_78556812_78563248_78607936_78609307_FR | 0.583598421 | 8.804321971 | 6.780547638 | 1.1202E-08 | 0.000118563 | 9.699358247 | 1.498582409 |
| CanFam3_20_8115423_8117184_8233713_8235237_FF | 0.427906461 | 8.250797621 | 6.264976766 | 7.3944E-08 | 0.000118563 | 7.932910688 | 1.345279983 |
| CanFam3_22_54934047_54938254_55122807_55123817_RF | 0.398321949 | 9.642706503 | 6.540143193 | 2.7035E-08 | 0.000118563 | 8.874946255 | 1.317974035 |
| CanFam3_6_39195094_39197108_39384777_39386801_RR | 0.392977649 | 9.716095099 | 6.749381088 | 1.2558E-08 | 0.000118563 | 9.592454091 | 1.313100782 |
| CanFam3_7_32179313_32195049_32246003_32249215_RR | 0.387507353 | 8.588065675 | 5.277698056 | 2.6025E-06 | 0.000312431 | 4.599056949 | 1.308131298 |
| CanFam3_17_62506776_62508089_62598339_62599667_FR | 0.359909126 | 8.555062228 | 3.848742909 | 0.00032722 | 0.005469501 | 0.105536445 | 1.283345058 |

| | | | | | | | |
|---|---|---|---|---|---|---|---|
| CanFam3_X_26083259_26086590_26123721_26124737_FF | 0.35843299 | 11.86978995 | 5.362979391 | 1.9223E-06 | 0.000271243 | 4.882430654 | 1.282032637 |
| CanFam3_4_57573278_57574839_57608066_57610552_FR | 0.356124014 | 8.494442116 | 5.119140211 | 4.5566E-06 | 0.00041652 | 4.075479109 | 1.279982436 |
| CanFam3_29_36199944_36202021_36306390_36308317_RF | 0.353355779 | 9.927458301 | 5.755239342 | 4.7093E-07 | 0.0001512 | 6.199028647 | 1.277528768 |
| CanFam3_30_3326453_3330215_3375157_3382851_RR | 0.351206662 | 8.299679976 | 6.538123509 | 2.7235E-08 | 0.000118563 | 8.868023641 | 1.275627109 |
| CanFam3_3_74009892_74014979_74210345_74211426_RF | 0.346487214 | 9.041854915 | 4.298305756 | 7.5973E-05 | 0.002129953 | 1.454676573 | 1.271461003 |
| CanFam3_2_78987543_78989579_79065537_79067880_RR | 0.345257665 | 8.248777466 | 5.845756541 | 3.3953E-07 | 0.000136731 | 6.505390534 | 1.270377852 |
| CanFam3_19_291779_295526_50826_55555_RR | 0.342759489 | 10.15410319 | 3.810783942 | 0.00036895 | 0.005921276 | -0.004861163 | 1.268179964 |
| CanFam3_33_7903610_7904731_8011733_8013746_RF | 0.342740822 | 11.15894963 | 5.775463785 | 4.3776E-07 | 0.000147803 | 6.267408948 | 1.268163555 |
| CanFam3_3_49400913_49414618_49489819_49496827_FF | 0.339649894 | 8.796228272 | 5.665987322 | 6.4964E-07 | 0.000170578 | 5.897782751 | 1.265449464 |
| CanFam3_2_45773482_45776169_45851310_45852835_FF | 0.33727441 | 10.3838751 | 3.897179433 | 0.00028053 | 0.004935261 | 0.24725472 | 1.263367539 |
| CanFam3_34_24844175_24862711_24903043_24906397_FR | 0.336600436 | 9.669580553 | 6.013053983 | 1.851E-07 | 0.000125381 | 7.073567692 | 1.262777478 |
| CanFam3_1_78079605_78081000_78180617_78197763_FF | 0.328389594 | 11.31811318 | 7.220671315 | 2.2299E-09 | 0.000118563 | 11.20791568 | 1.255611018 |
| CanFam3_35_16338409_16342034_16450580_16451875_RF | 0.314668604 | 8.41564831 | 6.029417046 | 1.7441E-07 | 0.000124599 | 7.129260019 | 1.243725929 |
| CanFam3_16_50972874_50981868_51036639_51039332_RR | 0.313948064 | 9.618249379 | 5.220527389 | 3.1865E-06 | 0.000346559 | 4.409761089 | 1.243104917 |
| CanFam3_17_19495826_19496909_19536615_19551007_RF | 0.313881641 | 9.576083928 | 5.560686725 | 9.4842E-07 | 0.000198716 | 5.543553308 | 1.243047685 |
| CanFam3_7_30900333_30903294_30922078_30924211_RR | 0.31377103 | 11.20280072 | 6.947176741 | 6.0794E-09 | 0.000118563 | 10.27085331 | 1.242952385 |
| CanFam3_16_51036639_51039332_51234447_51235965_RF | 0.311586046 | 9.962356772 | 5.43129009 | 1.5069E-06 | 0.00024192 | 5.110228238 | 1.241071338 |
| CanFam3_17_2880365_2881836_3072154_3075362_FF | 0.311347675 | 8.783614608 | 6.177315811 | 1.0181E-07 | 0.000118563 | 7.633460583 | 1.240866298 |
| CanFam3_17_388249_389308_586925_587959_RF | 0.308845492 | 10.52823394 | 3.879010375 | 0.00029723 | 0.005129984 | 0.193984395 | 1.238716028 |
| CanFam3_37_24290385_24298094_24398439_24402490_FF | 0.308775152 | 9.90679626 | 5.574121995 | 9.0379E-07 | 0.000194883 | 5.588673326 | 1.238655635 |
| CanFam3_2_61234321_61235994_61317580_61319613_RR | 0.308691578 | 9.726470196 | 5.861322552 | 3.2093E-07 | 0.000134887 | 6.558154199 | 1.238583882 |
| CanFam3_23_42804113_42805447_42848018_42850343_RF | 0.308341215 | 8.440906875 | 4.937786508 | 8.5979E-06 | 0.000584354 | 3.482485893 | 1.238283126 |
| CanFam3_X_36170434_36180921_36230157_36241354_RF | 0.305724333 | 9.422681259 | 5.989230195 | 2.0183E-07 | 0.000126859 | 6.992518133 | 1.236039059 |
| CanFam3_5_2020176_2021944_2175070_2177718_RR | 0.305509231 | 9.889407626 | 4.671653393 | 2.1557E-05 | 0.000994039 | 2.625419088 | 1.235854782 |
| CanFam3_22_12343094_12348728_12548557_12551045_FF | 0.304485331 | 9.320332591 | 5.570875325 | 9.1438E-07 | 0.000195393 | 5.577767846 | 1.23497799 |
| CanFam3_19_170899_172730_50826_55555_RR | 0.30371163 | 10.41787919 | 4.021000744 | 0.00018852 | 0.003817156 | 0.613719004 | 1.234315863 |
| CanFam3_35_24479504_24485855_24589397_24590654_FR | 0.303699242 | 9.041982476 | 4.311822947 | 7.2635E-05 | 0.002071144 | 1.496334196 | 1.234305265 |
| CanFam3_17_2880365_2881836_3048742_3051998_FR | 0.303460441 | 8.186849976 | 5.539741309 | 1.0224E-06 | 0.000205154 | 5.47325804 | 1.234100974 |
| CanFam3_22_59290338_59294926_59421522_59424000_RR | 0.302208624 | 10.36913196 | 5.998973437 | 1.9481E-07 | 0.0001263 | 7.025659878 | 1.233030618 |
| CanFam3_10_41684319_41686096_41808415_41814438_RR | 0.299623554 | 10.43456276 | 6.174416056 | 1.0289E-07 | 0.000118563 | 7.623562047 | 1.23082321 |
| CanFam3_25_11678603_11679802_11745739_11747571_RF | 0.297141262 | 9.950709859 | 6.551067404 | 2.5975E-08 | 0.000118563 | 8.912391261 | 1.228707284 |
| CanFam3_3_23717961_23719134_23743536_23749411_FF | 0.296323394 | 11.7611223 | 6.155597698 | 1.1019E-07 | 0.000118563 | 7.559335881 | 1.228010924 |
| CanFam3_27_31299498_31309546_31428583_31446337_FR | 0.295912195 | 9.647841354 | 5.018202234 | 6.4932E-06 | 0.000504287 | 3.744607107 | 1.227660964 |
| CanFam3_24_16270568_16284125_16434124_16437647_FR | 0.294643096 | 9.964770277 | 6.547195846 | 2.6346E-08 | 0.000118563 | 8.89912041 | 1.226581499 |
| CanFam3_26_3643865_3646009_3661573_3664056_FR | 0.294554469 | 9.871004266 | 5.60453054 | 8.1031E-07 | 0.000186333 | 5.690878971 | 1.226506151 |
| CanFam3_7_30379348_30385385_30564866_30568362_FR | 0.29142444 | 11.26517163 | 6.992457685 | 5.149E-09 | 0.000118563 | 10.42610721 | 1.223848043 |
| CanFam3_9_29107989_29115207_29331975_29350035_FR | 0.289444282 | 10.58060072 | 5.964931264 | 2.2044E-07 | 0.000128276 | 6.909897043 | 1.222169414 |
| CanFam3_18_28676686_28679289_28790557_28796160_FF | 0.289441441 | 9.691939487 | 6.79029591 | 1.0808E-08 | 0.000118563 | 9.732795874 | 1.222167008 |
| CanFam3_1_24964255_24967096_25111592_25113887_RR | 0.288625655 | 9.230179187 | 5.731427771 | 5.1318E-07 | 0.000155689 | 6.118574597 | 1.221476117 |
| CanFam3_35_1983390_1986171_2110889_2114605_RR | 0.288285623 | 10.26918959 | 6.040363065 | 1.6761E-07 | 0.000124212 | 7.166526071 | 1.221188258 |
| CanFam3_10_20688781_20693702_20822512_20824485_FF | 0.288022168 | 8.958409386 | 5.286169525 | 2.5255E-06 | 0.000307932 | 4.627153382 | 1.220965274 |
| CanFam3_7_17337513_17343563_17434574_17443671_FR | 0.286386061 | 8.81611084 | 6.270426534 | 7.2488E-08 | 0.000118563 | 7.95153988 | 1.219581407 |
| CanFam3_12_1102823_1107221_1176177_1181928_FR | 0.285544113 | 8.946941318 | 6.186171092 | 9.8573E-08 | 0.000118563 | 7.663691688 | 1.218869874 |
| CanFam3_7_76007147_76010110_76037945_76041916_RR | -0.554236006 | 9.556671926 | -5.899232419 | 2.7975E-07 | 0.000132861 | 6.686747961 | -1.468390826 |
| CanFam3_10_1812558_1815460_2069933_2072548_FR | -0.543586648 | 12.00425269 | -5.511159874 | 1.1326E-06 | 0.000213997 | 5.377428286 | -1.457591699 |
| CanFam3_11_28820782_28836360_28892204_28894641_FF | -0.532233988 | 11.58424656 | -6.6401275 | 1.8743E-08 | 0.000118563 | 9.217742296 | -1.446166826 |
| CanFam3_14_26668126_26677827_26823620_26835271_FF | -0.494491123 | 10.67068561 | -4.968541667 | 7.7237E-06 | 0.000552113 | 3.582573968 | -1.408823738 |
| CanFam3_24_26054542_26056955_26149901_26155964_RR | -0.490226785 | 14.66988963 | -6.951714932 | 5.979E-09 | 0.000118563 | 10.28641466 | -1.404665666 |

| | | | | | | | |
|---|---|---|---|---|---|---|---|
| CanFam3_3_17557004_17560145_17726672_17739743_FF | -0.48554687 | 10.24411767 | -4.400183274 | 5.4081E-05 | 0.001724986 | 1.770066384 | -1.400116495 |
| CanFam3_22_58366370_58370039_58559520_58561820_FR | -0.446035994 | 8.880942435 | -4.587605027 | 2.8719E-05 | 0.001175471 | 2.358405892 | -1.362292022 |
| CanFam3_23_368713_370013_506703_515684_FF | -0.434204243 | 9.002207898 | -4.060183617 | 0.00016606 | 0.003518874 | 0.730894815 | -1.351165356 |
| CanFam3_7_75993617_75995020_76037945_76041916_FR | -0.416306602 | 10.71669465 | -6.649849892 | 1.8087E-08 | 0.000118563 | 9.251083398 | -1.334506749 |
| CanFam3_26_24980431_24985909_25096321_25098486_FR | -0.40250071 | 12.75110103 | -6.414078779 | 4.2884E-08 | 0.000118563 | 8.443046919 | -1.321797076 |
| CanFam3_6_60993892_60995056_61092768_61094114_FF | -0.396936161 | 10.26933861 | -5.870002774 | 3.11E-07 | 0.000134887 | 6.587586948 | -1.316708657 |
| CanFam3_X_100272034_100276912_100343655_100346025_RF | -0.396561514 | 10.4128491 | -5.487250938 | 1.2338E-06 | 0.000220816 | 5.29734914 | -1.316366771 |
| CanFam3_X_97455526_97464290_97491109_97495161_FF | -0.383217848 | 10.74571867 | -6.156916301 | 1.0966E-07 | 0.000118563 | 7.563835539 | -1.304247662 |
| CanFam3_27_24481319_24483384_24615118_24617671_FR | -0.383075132 | 8.051467706 | -4.908049327 | 9.5353E-06 | 0.000620084 | 3.385905024 | -1.304118649 |
| CanFam3_6_29038683_29050190_29240029_29247478_FF | -0.382114875 | 12.73903593 | -4.756463005 | 1.6112E-05 | 0.000837089 | 2.896701508 | -1.303250917 |
| CanFam3_X_97455526_97464290_97522005_97525583_FR | -0.370298318 | 10.80026339 | -5.961949412 | 2.2284E-07 | 0.000128435 | 6.899761367 | -1.292620089 |
| CanFam3_4_45062189_45074917_45285074_45286167_FR | -0.369600501 | 13.37740131 | -4.853163713 | 1.1536E-05 | 0.000691961 | 3.208162306 | -1.291995012 |
| CanFam3_6_29038683_29050190_29240029_29247478_FR | -0.366280027 | 12.49841449 | -4.629051481 | 2.4936E-05 | 0.001082138 | 2.489842721 | -1.289024806 |
| CanFam3_38_11911932_11914685_11936045_11942843_FR | -0.358276681 | 9.722267848 | -4.868332012 | 1.0945E-05 | 0.000671284 | 3.257215676 | -1.281893743 |
| CanFam3_12_30321871_30325010_30340512_30346131_RF | -0.351166161 | 10.59794367 | -5.513855437 | 1.1217E-06 | 0.000212853 | 5.386461503 | -1.275591298 |
| CanFam3_20_34565970_34577697_34647610_34655536_RF | -0.349599349 | 9.831284183 | -5.624551462 | 7.5406E-07 | 0.000180609 | 5.75823307 | -1.274206718 |
| CanFam3_16_50017103_50022909_50222455_50232315_RR | -0.349378116 | 13.08309179 | -5.296342788 | 2.436E-06 | 0.000302759 | 4.660909618 | -1.274011337 |
| CanFam3_8_61551333_61552396_61581136_61584711_FF | -0.348247722 | 9.824923122 | -3.988245726 | 0.00020954 | 0.004090636 | 0.516205006 | -1.273013503 |
| CanFam3_25_35061477_35064415_35103921_35110189_FF | -0.347695972 | 10.87375081 | -4.43301727 | 4.8438E-05 | 0.001612831 | 1.872395547 | -1.272526739 |
| CanFam3_3_37186132_37187732_37262325_37265378_FF | -0.346502389 | 13.77337335 | -6.234288838 | 8.2707E-08 | 0.000118563 | 7.828035566 | -1.271474377 |
| CanFam3_X_97455526_97464290_97495161_97505186_FF | -0.344854403 | 11.45634855 | -5.238861601 | 2.9864E-06 | 0.000335357 | 4.470406113 | -1.270022805 |
| CanFam3_8_25127056_25135825_25262798_25272092_FR | -0.339821422 | 11.11279118 | -5.15263559 | 4.0496E-06 | 0.000391307 | 4.185708822 | -1.265599927 |
| CanFam3_23_4820027_4829464_4997994_5006043_RF | -0.339468944 | 11.26320855 | -5.002335585 | 6.8638E-06 | 0.000518519 | 3.692781653 | -1.265290755 |
| CanFam3_26_29429704_29434271_29528197_29529624_FR | -0.339189517 | 10.78129811 | -5.698761404 | 5.7731E-07 | 0.000162898 | 6.008301043 | -1.265045712 |
| CanFam3_X_97260653_97263737_97455526_97464290_RF | -0.338154631 | 10.7805167 | -5.52977442 | 1.0596E-06 | 0.00020826 | 5.439828087 | -1.264138584 |
| CanFam3_27_41239438_41243713_41337051_41350255_FR | -0.337883577 | 9.049240452 | -4.525623064 | 3.5442E-05 | 0.001331161 | 2.162722994 | -1.2639011 |
| CanFam3_X_97455526_97464290_97479084_97483786_FR | -0.33782138 | 10.86142091 | -5.731942453 | 5.1222E-07 | 0.000155685 | 6.120312959 | -1.263846612 |
| CanFam3_X_99789019_99793387_99995839_100001663_RF | -0.335522076 | 11.92300898 | -5.586158382 | 8.6558E-07 | 0.000191131 | 5.629114839 | -1.261833953 |
| CanFam3_X_97455526_97464290_97563032_97569353_FF | -0.334605541 | 10.9380962 | -5.514934299 | 1.1174E-06 | 0.000212633 | 5.390077194 | -1.261032572 |
| CanFam3_19_28225750_28227588_28262797_28268878_FF | -0.33370064 | 8.180995815 | -4.358521787 | 6.2168E-05 | 0.001882594 | 1.640699237 | -1.260241864 |
| CanFam3_3_48230774_48237747_48353947_48356862_RF | -0.330402844 | 9.647425135 | -5.109194534 | 4.7188E-06 | 0.000423571 | 4.042789506 | -1.257364419 |
| CanFam3_12_19071887_19075640_19273874_19280267_RR | -0.32852653 | 10.2181087 | -6.775006104 | 1.1432E-08 | 0.000118563 | 9.680350188 | -1.255730202 |
| CanFam3_21_7388705_7394830_7487331_7498278_FR | -0.325755575 | 10.23349359 | -5.133703991 | 4.3289E-06 | 0.00040519 | 4.123381237 | -1.253320661 |
| CanFam3_24_45842262_45845524_45993548_45998245_RR | -0.323888665 | 10.54142451 | -4.984036673 | 7.317E-06 | 0.000535969 | 3.633076186 | -1.25169986 |
| CanFam3_6_61060170_61061925_61092768_61094114_FF | -0.322709227 | 9.712144791 | -5.579667995 | 8.8598E-07 | 0.000193058 | 5.607305278 | -1.250676982 |
| CanFam3_19_39178236_39180425_39380979_39396482_FR | -0.321754935 | 11.67319908 | -4.261219985 | 8.5913E-05 | 0.002302819 | 1.340688696 | -1.249849977 |
| CanFam3_1_78180617_78197763_78375661_78378909_FF | -0.321307857 | 11.41091388 | -5.535697977 | 1.0373E-06 | 0.000206445 | 5.459694724 | -1.24946272 |
| CanFam3_13_39747682_39749517_39860646_39862238_RR | -0.319291407 | 9.984710267 | -6.262397879 | 7.4644E-08 | 0.000118563 | 7.924095653 | -1.24771757 |
| CanFam3_X_97455526_97464290_97495161_97505186_FR | -0.318167169 | 10.81384068 | -5.679664282 | 6.1843E-07 | 0.000167912 | 5.94388841 | -1.24674565 |
| CanFam3_4_30776968_30778661_30959175_30962265_RR | -0.317799738 | 9.895205324 | -4.638023801 | 2.4184E-05 | 0.001063657 | 2.518356609 | -1.246428164 |
| CanFam3_26_25238753_25241533_25464165_25468726_RR | -0.317579764 | 10.93053069 | -3.799203554 | 0.00038267 | 0.006068206 | -0.038422767 | -1.24623813 |
| CanFam3_13_50904752_50908493_51024982_51032173_FR | -0.316836007 | 13.56260907 | -4.008694788 | 0.00019617 | 0.003918159 | 0.577035974 | -1.245595819 |
| CanFam3_6_58288195_58302937_58462447_58468733_FF | -0.31584982 | 8.087831621 | -5.288345432 | 2.5061E-06 | 0.000306823 | 4.634371901 | -1.244744655 |
| CanFam3_4_47177458_47183642_47325716_47337092_FF | -0.314647111 | 10.27816138 | -3.909776693 | 0.00026948 | 0.004809073 | 0.284265913 | -1.243707401 |
| CanFam3_X_97455526_97464290_97563032_97569353_FR | -0.313485407 | 10.78258742 | -5.478363339 | 1.2736E-06 | 0.000224372 | 5.267601532 | -1.242706331 |

**Top Markers for DLBCL**

| 3D array probe | logFC | Average Expression | Moderated t-test for LIMMA | P Value | Adjusted P Value | B-statistics (log-odds) for LIMMA | Fold Change (FC) |
|---|---|---|---|---|---|---|---|
| CanFam3_27_25876660_25884084_26049678_26052242_FF | -0.47471111 | 8.382549015 | -2.487615334 | 0.01823899 | 0.172931299 | -3.201065916 | -1.389639926 |

| | | | | | | | |
|---|---|---|---|---|---|---|---|
| CanFam3_18_18520288_18523561_18639084_18642710_RR | -0.402318465 | 8.213489896 | -1.78165937 | 0.08426165 | 0.326988928 | -4.438714579 | -1.321630114 |
| CanFam3_X_101847546_101850983_101876697_101892356_RR | -0.491083457 | 7.363954094 | -3.849545008 | 0.00053103 | 0.080247921 | -0.173327479 | -1.405500003 |
| CanFam3_26_26176610_26178598_26237943_26244530_FF | -0.42080112 | 9.913204241 | -3.085708967 | 0.00415876 | 0.104226685 | -1.950355366 | -1.338670704 |
| CanFam3_18_18498471_18500877_18520288_18523561_RF | -0.365578742 | 8.206970738 | -1.568283224 | 0.12661413 | 0.392896484 | -4.749110108 | -1.288398372 |
| CanFam3_X_671335_674501_787107_788384_RR | -0.463687887 | 8.221762576 | -3.313475891 | 0.00228911 | 0.090789236 | -1.437583354 | -1.379062543 |
| CanFam3_3_25122972_25124688_25148378_25155983_FF | -0.467362747 | 8.033869953 | -3.614357016 | 0.00101672 | 0.081796666 | -0.736530282 | -1.382579795 |
| CanFam3_27_25757498_25766721_25876660_25884084_RF | -0.435248437 | 8.712888035 | -2.582818365 | 0.01455748 | 0.1585357 | -3.012729142 | -1.352143656 |
| CanFam3_13_61962433_61965230_62047816_62052621_RF | -0.421291631 | 8.035445006 | -1.996030388 | 0.05446653 | 0.270328482 | -4.095187667 | -1.339125925 |
| CanFam3_7_50664949_50670399_50728756_50730391_FR | -0.384307256 | 7.38222061 | -2.323215954 | 0.02665505 | 0.200876806 | -3.515307023 | -1.305232898 |
| CanFam3_17_12477211_12490460_12503850_12510011_RF | -0.438565063 | 7.4745818 | -4.849128885 | 3.0529E-05 | 0.080247921 | 2.308314734 | -1.355255689 |
| CanFam3_5_3085000_3086083_3099853_3105931_FR | -0.452584769 | 10.80803339 | -4.55057508 | 7.2463E-05 | 0.080247921 | 1.557822916 | -1.368489883 |
| CanFam3_13_62047816_62052621_62165633_62171199_FR | -0.416448098 | 8.079790929 | -1.887106415 | 0.06821073 | 0.298050755 | -4.273553566 | -1.334637641 |
| CanFam3_3_25122972_25124688_25330425_25335408_FR | -0.407090466 | 8.035505193 | -3.050184395 | 0.00455772 | 0.106912545 | -2.028734537 | -1.326008907 |
| CanFam3_8_8018241_8025786_8153862_8158547_FF | 0.640275179 | 10.04488046 | 3.493752335 | 0.00141162 | 0.084249539 | -1.020440564 | 1.558626423 |
| CanFam3_12_46277167_46279504_46401814_46408159_RR | 0.62409996 | 11.49390056 | 3.809349492 | 0.00059387 | 0.080247921 | -0.270393011 | 1.541249001 |
| CanFam3_3_42011223_42012378_42100327_42101824_RF | 0.574649462 | 7.994725743 | 6.599667712 | 1.8999E-07 | 0.078676958 | 6.660953208 | 1.489315554 |

**Top Markers for T-Zone Lymphoma**

| 3D array probe | logFC | Average Expression | Moderated t-test for LIMMA | P Value | Adjusted P Value | B-statistics (log-odds) for LIMMA | Fold Change (FC) |
|---|---|---|---|---|---|---|---|
| CanFam3_33_3068933_3071158_3158267_3159448_FR | -0.482770241 | 7.615821327 | -4.390871394 | 0.00011758 | 0.046367907 | 1.164838283 | -1.397424405 |
| CanFam3_7_69415658_69417041_69558041_69559499_RF | 0.411048521 | 8.404551566 | 2.842180926 | 0.00777581 | 0.088284581 | -2.509232155 | 1.329651827 |
| CanFam3_1_26894587_26898428_26993061_26995825_FR | 0.478372102 | 7.818399063 | 3.905891335 | 0.00046186 | 0.046367907 | -0.042956531 | 1.393170763 |
| CanFam3_29_20063845_20066056_20103813_20106226_RR | 0.471038011 | 6.826751544 | 3.771544747 | 0.00067019 | 0.04660978 | -0.370947713 | 1.386106405 |
| CanFam3_15_22109294_22110421_22182809_22186420_RR | 0.449777301 | 6.594636722 | 5.595315577 | 3.6361E-06 | 0.046367907 | 4.226751167 | 1.365829406 |
| CanFam3_15_41400908_41406654_41517768_41525940_RF | -0.462661201 | 7.429707489 | -4.603190418 | 6.403E-05 | 0.046367907 | 1.701705367 | -1.378081489 |
| CanFam3_35_3482632_3484835_3499602_3501391_RR | -0.396786049 | 7.471383712 | -2.082763659 | 0.04543572 | 0.196223439 | -3.994080893 | -1.316571661 |

**Top Markers for Hemangiosarcoma**

| 3D array probe | logFC | Average Expression | Moderated t-test for LIMMA | P Value | Adjusted P Value | B-statistics (log-odds) for LIMMA | Fold Change (FC) |
|---|---|---|---|---|---|---|---|
| CanFam3_15_18744079_18746522_18763203_18764950_FR | -0.374935552 | 7.507867913 | -1.397746383 | 0.17186655 | 0.412767134 | -5.061771312 | -1.296781623 |
| CanFam3_24_39311015_39312144_39499938_39502137_FF | -0.453677454 | 6.979741879 | -3.279675448 | 0.00252278 | 0.045328044 | -1.544597491 | -1.369526758 |
| CanFam3_9_32111009_32117007_32291966_32295647_RF | -0.926437929 | 12.6908453 | -2.109670578 | 0.04285183 | 0.190848559 | -3.972170378 | -1.900577597 |
| CanFam3_23_35495556_35497783_35635274_35651194_FF | -0.462443861 | 8.290741525 | -4.26474607 | 0.00016726 | 0.029090383 | 0.859577115 | -1.377873899 |
| CanFam3_25_48634487_48635577_48859327_48866621_FR | -0.475350893 | 8.151048402 | -4.642257442 | 5.6768E-05 | 0.029090383 | 1.821750847 | -1.390256318 |
| CanFam3_6_50983197_50984903_51123223_51125507_RR | -0.426782912 | 7.381858318 | -4.372274815 | 0.00012313 | 0.029090383 | 1.132278481 | -1.344232707 |
| CanFam3_19_19668047_19671316_19883962_19885557_FF | 0.313933934 | 7.676268851 | 2.115848964 | 0.04228447 | 0.189438561 | -3.961207045 | 1.243092742 |
| CanFam3_25_48541987_48544892_48634487_48635577_RF | -0.440909646 | 8.109643156 | -4.462908304 | 9.5019E-05 | 0.029090383 | 1.363058712 | -1.357459962 |
| CanFam3_14_9326366_9339564_9546090_9552599_RF | -0.379124861 | 7.937606007 | -2.661145196 | 0.01210499 | 0.097172392 | -2.90788657 | -1.300552701 |

**Top Markers for Histiocytic Sarcoma**

| 3D array probe | logFC | Average Expression | Moderated t-test for LIMMA | P Value | Adjusted P Value | B-statistics (log-odds) for LIMMA | Fold Change (FC) |
|---|---|---|---|---|---|---|---|
| CanFam3_1_8727830_8732342_8830326_8834568_RF | -0.455724785 | 7.839690666 | -2.325421996 | 0.02654619 | 0.140541663 | -3.619350534 | -1.371471636 |
| CanFam3_34_23862723_23865328_24147458_24148561_RF | -0.460041667 | 7.459753851 | -2.440453319 | 0.02038455 | 0.121697585 | -3.394782097 | -1.375581546 |
| CanFam3_2_84003200_84004536_84054773_84060215_RF | 0.419759075 | 7.757681794 | 2.775040264 | 0.00913853 | 0.080011604 | -2.702318651 | 1.337704144 |
| CanFam3_27_29299062_29303517_29331293_29345286_RR | 0.413082404 | 6.587020612 | 2.815472835 | 0.00826781 | 0.076078403 | -2.615004622 | 1.331527665 |
| CanFam3_10_14732268_14733431_14833087_14836743_RF | -0.422605835 | 6.513209558 | -3.568576874 | 0.00115598 | 0.032228683 | -0.873695416 | -1.34034634 |
| CanFam3_22_56698814_56701176_56824010_56827219_FF | -0.346222944 | 7.124936029 | -2.213947007 | 0.03407723 | 0.16124228 | -3.829762763 | -1.271228121 |
| CanFam3_22_33574242_33576183_33596207_33601110_FF | -0.421533822 | 7.509313471 | -4.77112026 | 3.8584E-05 | 0.017681346 | 2.186331002 | -1.339350749 |
| CanFam3_24_11225946_11232495_11346764_11358311_FF | -0.450137228 | 13.68419452 | -3.467933877 | 0.00151834 | 0.035837576 | -1.117212152 | -1.3661702 |
| CanFam3_13_21806403_21809364_22013299_22016769_RR | 0.377784082 | 7.673849885 | 2.925480812 | 0.00627609 | 0.066497872 | -2.373820653 | 1.299344585 |
| CanFam3_38_10911611_10913415_11079662_11095682_FR | -0.429041225 | 6.552541231 | -5.536177797 | 4.1698E-06 | 0.017681346 | 4.191070146 | -1.34633854 |

| | | | | | | | |
|---|---|---|---|---|---|---|---|
| CanFam3_23_42374336_42380491_42415529_42419010_RF | -0.409191776 | 10.67949004 | -4.131913117 | 0.00024133 | 0.020169787 | 0.532802871 | -1.327941668 |
| CanFam3_5_88590564_88592906_88638120_88639730_FF | 0.354173888 | 6.399125973 | 2.887703106 | 0.00690271 | 0.069679543 | -2.457229521 | 1.278253422 |
| CanFam3_9_3331047_3332297_3370937_3374814_FF | -0.345016479 | 6.875141928 | -2.17429174 | 0.03718651 | 0.169199118 | -3.90283089 | -1.27016549 |
| CanFam3_21_50252490_50255889_50489194_50490728_FF | -0.401013049 | 8.498047409 | -3.930161793 | 0.00042585 | 0.023117853 | 0.021756396 | -1.320434784 |
| CanFam3_X_68172047_68186818_68258559_68259788_FF | -0.379916503 | 11.01958127 | -4.604108214 | 6.2525E-05 | 0.017681346 | 1.750712031 | -1.301266542 |
| **Top Markers for Osteosarcoma** | | | | | | | |
| **3D array probe** | **logFC** | **Average Expression** | **Moderated t-test for LIMMA** | **P Value** | **Adjusted P Value** | **B-statistics (log-odds) for LIMMA** | **Fold Change (FC)** |
| CanFam3_37_2363419_2371539_2496561_2498045_RR | -0.472156339 | 7.950119192 | -2.245887388 | 0.03176687 | 0.233172092 | -3.659993512 | -1.387181284 |
| CanFam3_21_10332790_10336978_10454491_10457037_RR | 0.335104439 | 7.782924597 | 1.328945888 | 0.19329097 | 0.526852013 | -5.058768737 | 1.261468725 |
| CanFam3_X_87179887_87185947_87244440_87245978_FF | -0.414722025 | 13.57598922 | -1.525768741 | 0.13691935 | 0.452545184 | -4.808786368 | -1.333041805 |
| CanFam3_5_58469072_58472778_58602674_58604717_FR | 0.431579563 | 7.550576532 | 2.647159193 | 0.01250357 | 0.154862419 | -2.885322338 | 1.34870943 |
| CanFam3_13_7501436_7503370_7524195_7527457_RR | 0.389796726 | 7.827771069 | 2.712626466 | 0.01066593 | 0.144810767 | -2.751379614 | 1.310208784 |
| CanFam3_32_38115147_38120145_38308888_38324226_FF | -0.421734507 | 9.479075223 | -3.17532814 | 0.00330918 | 0.093770215 | -1.754031333 | -1.339537071 |
| CanFam3_22_56356278_56362583_56426305_56427322_FR | -0.40290895 | 6.668859886 | -3.359924674 | 0.00203348 | 0.081913912 | -1.334868876 | -1.322171159 |
| CanFam3_5_50175581_50178428_50208523_50212894_FR | -0.421847462 | 7.784939858 | -3.071737543 | 0.00432893 | 0.102614759 | -1.984374701 | -1.339641953 |
| CanFam3_16_20752908_20754401_20777017_20781577_FR | -0.417667596 | 7.680677805 | -3.909047405 | 0.00045348 | 0.072674267 | -0.034873278 | -1.335766276 |
| CanFam3_15_40284024_40286232_40298694_40305948_FR | 0.375295361 | 7.792613567 | 3.293912266 | 0.0024231 | 0.085549154 | -1.485974054 | 1.297105082 |
| CanFam3_32_38126234_38135766_38308888_38324226_FF | -0.374614095 | 9.448421973 | -3.007493517 | 0.00510441 | 0.108836873 | -2.125313384 | -1.296492711 |
| CanFam3_16_38985349_38986956_39018382_39025503_FR | 0.394163054 | 7.882377985 | 3.265320751 | 0.00261321 | 0.08731584 | -1.551012199 | 1.314180148 |
| CanFam3_1_50579723_50581242_50642045_50647459_RR | -0.41125037 | 6.891813377 | -4.301398276 | 0.00014982 | 0.072674267 | 0.928284086 | -1.329837873 |
| **Top Markers for Melanoma** | | | | | | | |
| **3D array probe** | **logFC** | **Average Expression** | **Moderated t-test for LIMMA** | **P Value** | **Adjusted P Value** | **B-statistics (log-odds) for LIMMA** | **Fold Change (FC)** |
| CanFam3_16_24898866_24901480_24986987_24992147_RR | -0.487299676 | 8.202436962 | -2.27542605 | 0.02976283 | 0.152851914 | -3.739020475 | -1.401818604 |
| CanFam3_5_37031042_37040176_37109754_37116066_RF | -0.499430749 | 6.720708772 | -3.16580486 | 0.00340042 | 0.058797577 | -1.847823333 | -1.413655659 |
| CanFam3_5_43083373_43094213_43318874_43328027_FF | -0.545708324 | 8.985725419 | -5.682425395 | 2.787E-06 | 0.006138661 | 4.576902214 | -1.459736859 |
| CanFam3_4_19313435_19315467_19418384_19421902_FF | -0.548679294 | 9.199796381 | -5.473545988 | 5.1076E-06 | 0.007153924 | 4.028818761 | -1.462746022 |
| CanFam3_22_5180335_5183126_5377750_5378964_RF | -0.522243068 | 12.96891473 | -7.734835454 | 8.4349E-09 | 0.000653651 | 9.75382037 | -1.436186461 |
| CanFam3_32_4753227_4757060_4907861_4910959_RR | -0.516593547 | 10.97937426 | -6.491376013 | 2.7169E-07 | 0.002807205 | 6.672758109 | -1.430573426 |
| CanFam3_34_41554876_41558955_41716884_41721444_RF | 0.641626133 | 7.035042457 | 5.246593174 | 9.8705E-06 | 0.008428654 | 3.431846075 | 1.56008662 |
| CanFam3_10_65724007_65725431_65882395_65885162_RR | 0.594467287 | 8.214038025 | 3.909373903 | 0.00045507 | 0.028041177 | -0.04249187 | 1.509914949 |

RP/RSum – Rank sum,
FC:(class1/class2) – average computed fold change,
pfp – percentage false positive,
t- moderated t-test for LIMMA,
B – B-statistics (log-odds)
